# HCM-associated mutations in MYH6/7 drive pathologic expression of TGF-β1 in cardiomyocytes within weeks of developmental specification

**DOI:** 10.1101/2024.08.08.606705

**Authors:** Jeanne Hsieh, Megan A. L. Hall, Mohammad Shameem, Patrick J. Ernst, Forum Kamdar, Bhairab N. Singh, Robert L. Meisel, Brenda M. Ogle

## Abstract

Hypertrophic cardiomyopathy (HCM) is characterized by myocyte hypertrophy, sarcomere disarray, and myocardial fibrosis, leading to significant morbidity and mortality. As the most common inherited cardiomyopathy, HCM largely results from mutations in sarcomeric protein genes. Current treatments for HCM primarily focus on alleviating late-stage symptoms, with a critical gap in the detailed understanding of early-stage deficiencies that drive disease progression. We recently showed, in monolayers of cardiomyocytes derived from human induced pluripotent stem cells (hiPSCs) with *MYH7* R723C and *MYH6* R725C mutations, altered expression of several extracellular matrix (ECM)-related genes with associated defects in cardiomyocyte-ECM adhesion. To better evaluate the cardiomyocyte-ECM interface and pathological ECM dynamics in early-stage HCM, here we adopted a 3D engineered heart tissue (EHT) model containing both cardiomyocytes and fibroblasts, the primary contributor to ECM remodeling. Mutant EHTs showed aberrant cardiomyocyte distribution, augmented calcium handling, and force generation compared to controls. Altered proteoglycan deposition and increased phosphorylated focal adhesion kinase (pFAK) further indicated changes in ECM composition and connectivity. Elevated transforming growth factor beta-1 (TGF-β1) secretion and a higher proportion of activated fibroblasts were identified in mutant EHTs, along with sustained TGF-β1 transcription specifically in mutant cardiomyocytes. Remarkably, blocking TGF-β1 receptor signaling reduced fibroblast activation and contraction force to control levels. This study underscores the early interplay of mutant hiPSC-CMs with fibroblasts, wherein mutant cardiomyocytes initiate fibroblast activation via TGF-β1 overexpression, independent of the immune system. These findings provide a promising foundation for developing and implementing novel strategies to treat HCM well before the manifestation of clinically detectable fibrosis and cardiac dysfunction.

## Introduction

Hypertrophic cardiomyopathy (HCM) stands as the predominant heritable cardiomyopathy, affecting approximately 1 in 200-500 individuals globally [1, 2, 3]. The majority of HCM-causing mutations exist in gene loci encoding sarcomeric proteins, especially myosin heavy chain (MHC) and myosin binding protein C (MyBP-C), leading to immediate alterations in cardiomyocyte (CM) mechanical function. Even so, the symptomatic clinical manifestations of HCM are not typically observed until well after birth which is when most historical studies of the mechanisms of HCM have been conducted. We and others have recently used hiPSC-CM monolayer to define early-stage, phenotypic changes at the cellular scale [4, 5]. In particular, we identified changes in ECM dynamics, integrin expression and cellular adhesion within weeks of differentiation of mutant CMs [4]. Building upon this observation, here we sought a multi-cellular, human model to delve deeper into myocyte functional alterations associated with early-stage ECM remodeling and elucidate associated mechanisms underlying early-stage disease progression.

One significant challenge in studying human HCM is to obtain early-stage disease-specific cardiac tissue samples [6]. Over recent decades, cardiac tissue engineering using gene-edited human induced pluripotent stem cells (hiPSCs) has presented a promising avenue for creating *in vitro* tissue models to explore human cardiac functional changes induced by HCM mutations [7–10]. As mutations in the *MYH7* locus, particularly within the converter domain of β-MHC, can lead to severe HCM phenotypes in patients [11], we focused on evaluating the impact of β-MHC mutations on functional phenotypes within the context of engineered heart tissues (EHTs). Although the majority of HCM patients harbor heterozygous HCM mutations, protective mechanisms such as allelic imbalance [12–14] often delay disease progression until later in life. To accelerate the effects directly associated with HCM mutations at early time points, we utilized CRISPR-Cas 9-mediated base editing to generate homozygous HCM MHC mutation onto a healthy hiPSC line, namely *MYH7* c.2167C > T (R723C) which also contained a heterozygous *MYH6* c.2173C > T (R725C) mutation [4].

The human heart is an intricate and multi-cellular organ. In addition to cardiomyocytes, various supporting cells including fibroblasts, endothelial cells, and smooth muscle cells play vital roles in upholding overall heart function. These supporting cells contribute to the structural integrity of the heart, regulate ECM dynamics, facilitate vascularization, and regulate signaling pathways crucial for cardiac homeostasis. Moreover, the interplay among various cell types within the cardiac microenvironment exerts profound effects on diverse physiological processes, such as electrical signaling, contractility, remodeling, and responses to pathological stimuli. These interactions not only influence cardiac function but have also been shown to play a significant role in the onset and progression of certain cardiac diseases [15, 16].

Engineered heart tissues (EHTs) are advanced three-dimensional *in vitro* models consisting of cardiomyocytes and other supportive cells embedded in a biomaterial matrix. EHTs have been utilized to mimic some of the structural and functional characteristics of native cardiac tissue, facilitating the study of cardiac physiology, pathophysiology, and drug responses. One key advantage of EHTs is their ability to elucidate the role of support cells, such as fibroblasts and endothelial cells, in enhancing the molecular maturation and function of both healthy and diseased hiPSC-CMs [17–20]. Through paracrine signals, ECM production, or direct cell-cell interaction mediated by fibroblasts, incorporating them into engineered tissues has been shown to improve electrophysiological properties [20–28]. Moreover, cardiomyocytes co-cultured with fibroblasts exhibit increased synchronous tissue contractions compared to those comprised solely of cardiomyocytes [23, 25, 26]. Understanding the complexities of intercellular communications is essential for unraveling the mechanisms underlying cardiac diseases and developing targeted therapeutic strategies to mitigate their progression [17, 19].

In this study, we utilized an engineered 3D cardiac tissue model, incorporated with supporting cells, to better recapitulate the pathological changes associated with HCM mutations. Considering fibroblasts are the main source of ECM remodeling [29, 30], healthy fibroblasts were incorporated into the EHTs, further improving the ability of this *in vitro* tissue to mimic *in vivo* cardiac tissues. To investigate the functional pathology of MHC mutant hiPSC-CMs in a 3D system, we designed a PDMS culture well that contains two fixed posts to secure the EHTs and provide passive strain during cultivation. This static, passive strain of the posts models afterload, the pressure the heart muscle must work against to pump blood [31]. Using this model, we find elevated force of contraction, altered calcium handling, and changed ECM dynamics in the mutant EHTs. Further, we find TGF-β1 signaling upregulated with disease with differential expression in CM, suggesting CMs are the primary mediator of fibroblast activation within 30 days of EHT generation and 45 days after cardiomyocyte specification. This study offers an engineered 3D model for studying early-stage HCM and identifies a potential therapeutic strategy to mitigate the progression of this debilitating cardiac disease well before the onset of clinical symptoms.

## Material and methods

### 1. hiPSC lines maintenance and cardiac differentiation

The control hiPSC line was kindly provided by Dr. Jianyi Zhang (University of Alabama at Birmingham), this hiPSC line was reprogrammed from cardiac fibroblasts of a healthy female donor and modified to overexpress cyclin D2 to enhance the yield of cardiac differentiation [32, 33]. The *MYH7* and *MYH6* mutations were edited onto the control hiPSC line via CRISPR/Cas9 as published previously [4, 34].

To obtain *MYH7/MYH6* mutant and wild-type control hiPSC-CMs for fabricating EHTs, a fully defined direct cardiac differentiation by modulating Wnt/β-catenin signaling was performed. Briefly, both mutant and control hiPSC lines were cultured on Matrigel-coated 6-well plates and fed with mTeSR1 medium for 5 days to reach 80%–90% confluency. All hiPSC lines used for differentiation were between passages 57–78. To harvest the hiPSCs, culture medium was removed, each well was washed one time with sterile DPBS before adding 1 mL of room-temperature Accutase. The plates were transferred into the incubator to allow Accutase to react at 37°C for 8 minutes. 0.5 mL of mTeSR1 medium was added into each well to stop Accutase reactions. The singularized cells were collected, centrifuged in 200g for 5 minutes to Accutase solution, and resuspended in mTeSR1 with 5μM ROCK inhibitor (Y27632). The hiPSCs were seeded with the density of 1 × 10^6^ cells/well on Matrigel-coated 12-well plates and maintained in the 5% CO_2_ incubator at 37°C for 24 hours. After 24 hours, on Day -1, 1 mL of fresh mTeSR1 medium was added to each well. On Day 0, initiation of cardiac differentiation was performed on 100% confluent hiPSCs by adding 8μM GSK3-β inhibitor (CHIR99021) in 2 mL of RPMI/B-27 without insulin (RPMI minus) medium into each well for 24 hours in the 37°C, 5% CO2 incubator to induce mesoderm. On Day 1, CHIR99021 medium was replaced with fresh RPMI/B27 minus medium. On Day 3, 5μM Wnt inhibitor (IWP2) in 2 mL of RPMI minus medium was added into each well to induce cardiac differentiation. On Day 5, fresh RPMI/B27 minus insulin was added into each well. On Day 7 and every 3 days thereafter, fresh maintenance medium, RPMI/B-27 medium, was added to each well. Spontaneous twitching was observed on Day 8, and robust spontaneous contraction occurred by Day 12. To purify the cardiomyocyte population, cardiomyocyte enrichment was performed on Day 10–14 by adding 2 mL DMEM (without glucose) with 4 mM sodium L-lactate every 2 days. On Day 14–16, cells were maintained in RPMI/B27 medium. Day 16 hiPSC-CMs were harvested by treating the wells with 0.25% Trypsin-EDTA for 12 minutes at 37°C. The singularized hiPSC-CMs were resuspended in RPMI/B-27 medium with 20% fetal bovine serum and 5μM ROCK inhibitor (Y27632) for recovery for 30 minutes before being used for fabricating EHTs.

#### 1. 2. EHT fabrication

For EHT formation and maintenance, PDMS culture wells were utilized in this study. A customized 3D-printed negative template was used to fabricate the PDMS wells. The Sylgard 184 PDMS precursor and the curing agent were mixed at the 10:1 mass ratio, and vacuumed for 1 hour at room temperature before being cured at 48°C for 24 hours on the negative template. The molded PDMS wells were carefully removed from the template and cured at 120°C for 7 days. After the curing process, PDMS wells were sonicated in 70% ethanol for 30 minutes, autoclaved, treated with oxygen plasma (PDC-32G, Harrick Plasma), and coated with 0.5% Pluronic F-127 solution. Before fabricating EHTs, the PDMS wells were washed with sterile DPBS to remove the Pluronic solution and exposed to 254 nm ultraviolet light for 30 minutes in a biological safety cabinet. Each PDMS well has the dimension of 15 mm in length, 5 mm in width and depth, and contains two rectangular posts (3.2 mm in width, 0.8 mm in thickness, and 0.45 mm in height) positioned 1 mm from the short end walls of the well. To fabricate an EHT, 1.6 × 10^6^ hiPSC-CMs and 0.4 × 10^6^ human dermal fibroblasts (Lonza, Cat. # CC-2509) were mixed in 60 μL of RPMI/B27 medium with 20μM Aprotinin and 5μM ROCK inhibitor, then adding 120 μL of 20mg/mL fibrinogen to the cell mixture before adding 20 μL of 100U/mL thrombin to solidify EHT in a PDMS culture well.

#### 1. 3. Calcium transient measurement

Movement of Calcium ions was assessed with DMi8 fluorescence microscope (Leica, Wetzlar, Germany) and LAS X software. EHTs were incubated with 5 μM Fluo-4 acetoxymethyl ester (Fluo-4 AM) in culture medium at 37°C for 30 minutes, and then culture medium was exchanged to Tyrode’s salt solution for another 30 minutes incubation at 37°C. After that, transfer the EHT plates onto the microscope stage and covered with a heating plate to maintain the temperature at 37°C. Fluo-4 AM intensity was recorded at the frame rate of 6.90 Hz with a 30 ms exposure time. The acquired data was processed by ImageJ to obtain a time trace of calcium signal and then analyzed in a custom-written Python script to extract maximal and minimal intensity and corresponding time points for each peak. Peak amplitude was determined by F/F_0_ where F = <F_max_> - <F_min_> and F_0_ = <F_min_>. <F_max_> and <F_min_> represent the averaged maximal and minimal intensity for each peak, respectively.

#### 1. 4. Force generation measurement

Contraction force of EHT was measured using Mach-1 equipment with a force transducer (GSO-25, capacity range: 25g). Before measuring force, EHTs were incubated in TSS buffer for 30 minutes at 37°C, and transferred to a 3D-printed well containing TSS buffer on a heating plate to maintain the system at 37°C. EHT was then held on two mounting needles, one mounting needle had a fixed location and the other was connected with the force transducer. Force generation measurement started at unstretched EHT length (normally around 12mm) for recording 30 seconds without pacing, and then applied 1 Hz pacing for another 30-second recording. And then the EHT was stretched for 5% of its initial length for non-pacing recording and followed by paced recording, repeat the 5% stretching and recording process until the length reached 120% of the initial length. After the non-pacing and 1 Hz pacing for 120% length, recording force for 2 Hz and 3 Hz pacing condition.

#### 1. 5. Immunohistochemical staining

Immunohistochemical staining was performed on frozen tissue sections. EHTs were firstly fixed with 4% paraformaldehyde (PFA) overnight, then washed and stored in PBS at 4°C. Before embedding tissues in OCT (Tissue-Tek®), the tissues were placed in 15% sucrose in PBS until they sank, and then transferred to 30% sucrose in PBS overnight or until they sank. Tissues were placed in cryo molds and covered with sufficient OCT to avoid bubbles or contact with the air surface interface. Embedded tissues were stored at -80°C prior to cryostat sectioning. Frozen tissues were cut at 4-10 µm and mounted on gelatin-coated histological slides. Cryostat temperature was maintained between -15 and -23°C. Frozen sections were air-dried for 30 minutes at room temperature to prevent sections from falling off the slides during staining. Sample slides were stored at -20°C prior to staining.

To perform immunostaining, sample slides were permeabilized by 0.2% Triton X-100 at room temperature for 1 hour, washed with PBS three times, and then incubated with blocking buffer (2.5 g non-fat dry milk in 50 mL of 0.2% Triton X-100) at room temperature for 2 hours. After washing with PBS three times, the primary antibody was added for overnight incubation at 4°C. The slides were then washed twice with 0.2% Tween 20 and twice with PBS before incubating with the secondary antibody (Alexa Fluor 488 goat anti-mouse IgG, concentration 1:500; Alexa Fluor 647 goat anti-rabbit IgG, concentration 1:500) for 1.5 hours at room temperature. After another series of washes with 0.2% Tween 20 and PBS, the slides were mounted with DAPI/DABCO. Images were taken using a Leica fluorescence microscope.

#### 1. 6. Protein isolation and Western blotting

EHTs were washed with 1x PBS and homogenized in the 1x RIPA lysis buffer containing protease and phosphatase inhibitors. Tissue lysates were centrifuged at 14,000*g* for 20 minutes at 4°C, and the supernatants were collected. Total protein was quantified using Bio-Rad protein assay dye reagent. For each sample, a total of 30 μg of protein was mixed with 1x SDS loading buffer, denatured at 99°C, loaded onto a 14% gradient SDS polyacrylamide gel, and separated electrophoretically. The electrophoresed proteins were transferred to PVDF membranes. After transfer, blots were incubated in the blocking buffer (5% skim milk in TBST) for 1 hour and then with primary antibodies overnight at 4°C. Primary antibodies were detected with appropriate horseradish peroxidase-coupled secondary antibodies using the iBright CL1500 imaging system (Invitrogen, MA, USA). Signal intensity was quantified using Image J software.

#### 1. 7. Focal adhesion complex analysis

For focal adhesion complex experiments, hiPSC-CMs were seeded in a Matrigel-coated 24-well plate. After 24 hours, the cells were transfected with the vinculin-GFP plasmid using Lipofectamine Stem Transfection Reagent (Thermo Fisher Scientific, Waltham, MA, USA) according to the manufacturer’s protocols. After 72 hours, GFP images were captured using a confocal microscope (Olympus IX83, Waltham, MA, USA).

#### 1. 8. RNAscope in situ hybridization

In situ hybridization was performed using formalin-fixed paraffin-embedded (FFPE) samples, following the protocols provided by Advanced Cell Diagnostics (Newark, CA, USA). The probe used in this procedure targets mRNA encoding Homo sapiens transforming growth factor, TGFB1 (RNAscope™ Probe-Hs-TGFB1-long, Cat. # 407121, accession number: NM_000660.4, target region: 170 – 1876 bp).

Briefly, 4-5 μm sections of FFPE samples were deparaffinized by submerging in xylene for 5 minutes twice and dehydrated by submerging in 100% ethanol for 2 minutes twice, followed by drying in an oven at 60°C for 5 minutes. To pretreat the samples, 75 μL of RNAscope Hydrogen Peroxide was used to cover each section and incubated at room temperature for 10 minutes. After washing the slides in distilled water, they were slowly submerged into distilled water at 98-102°C for 10 seconds and then immediately moved into 1x RNAscope Antigen Retrieval solution at 98-102°C for 15 minutes. Once the slides cooled down and dried, hydrophobic barriers for each section were drawn. 75 μL of RNAscope Protease Plus was added to cover each section and incubated at 40°C for 15 minutes. The RNAscope assay started with a 2-hour incubation of TGFB1 probe in the HybEZ Oven. The probe solution was diluted 1:1 with probe diluent (Advanced Cell Diagnostics), and each tissue section received 75 μL of the solution. After probe hybridization, AMP 1, AMP 2, and AMP 3 were applied to effectively and specifically amplify the targeted signals. Incubation for AMP 1 and AMP 2 was 30 minutes at 40°C and 15 minutes at 40°C for AMP 3. Between each step, slides were submerged into wash buffer for 2 minutes per rinse (8.7 g/L of sodium chloride, 4.41 g/L of sodium citrate, 3 g/L of sodium dodecyl sulfate; based on Wang et al. (2012)). HRP-C1 solution was then applied and incubated at 40°C for 15 minutes. For RNA signals, Opal 570 dye (Akoya Biosciences, Marlborough, MA, USA; concentration 1:750) was used and incubated at 40°C for 30 minutes. Before the immunohistochemistry (IHC) process, blocker solution was added and incubated at 40°C for 15 minutes. Primary antibodies containing 1:200 mouse anti-human cTnT and 1:200 rabbit anti-human αSMA diluent were applied and incubated at 4°C overnight. The next day, the slides were washed in 1x TBS buffer three times for 10 minutes per rinse before adding the secondary antibody (1:500 goat anti-mouse 647 and 1:500 goat anti-rabbit 488 diluent) solution and incubating at room temperature for 1 hour. After washing the slides in 1x TBS buffer three times for 10 minutes per rinse, 100 μL of ProLong Gold Antifade Mountant with DAPI counterstain solution was added for each section to cover and mount the slides with a coverslip.

#### 1. 9. Statistical analysis

All experiments were repeated at least two times and the values presented are mean ± standard error of the mean (SEM) from the replicates of at least two independent experiments. Statistical significance was determined using the Student’s t-test when comparing two groups and ONE-WAY ANOVA with multiple comparisons when comparing more than two groups. A p-value < 0.05 was considered a significant change and was highlighted in each panel by an asterisk.

## Results

### 1. EHT fabrication and macroscale appearance were consistent between wild-type and HCM mutant groups

Mutant hiPSC lines were generated through CRISPR/Cas9 editing of a wild-type hiPSC line, as detailed in our previously published studies [4]. Subsequently, cardiac differentiation and lactate purification were performed to obtain Day 16 hiPSC-CMs. In this study, we employed fibrin-based EHTs to examine the impact of HCM mutations on cardiac function at Day 15 and 30 after EHT fabrication as shown in Figure 1A. The inclusion of fibroblasts was crucial in this EHT model, as they served as the primary source of ECM deposition immediately following EHT fabrication, facilitating tissue organization. To explore the HCM pathology initiated by MHC mutations in hiPSC-CMs, diseased EHTs were fabricated by combining mutant hiPSC-CMs with wild-type fibroblasts, while control EHTs were constructed using isogenic control hiPSC-CMs and wild-type fibroblasts. Importantly, prior to integration into the EHTs, hiPSC-CMs exhibited spontaneous beating, and this rhythmic activity resumed within two days of EHT fabrication for both mutant and control groups.

**Figure 1.**
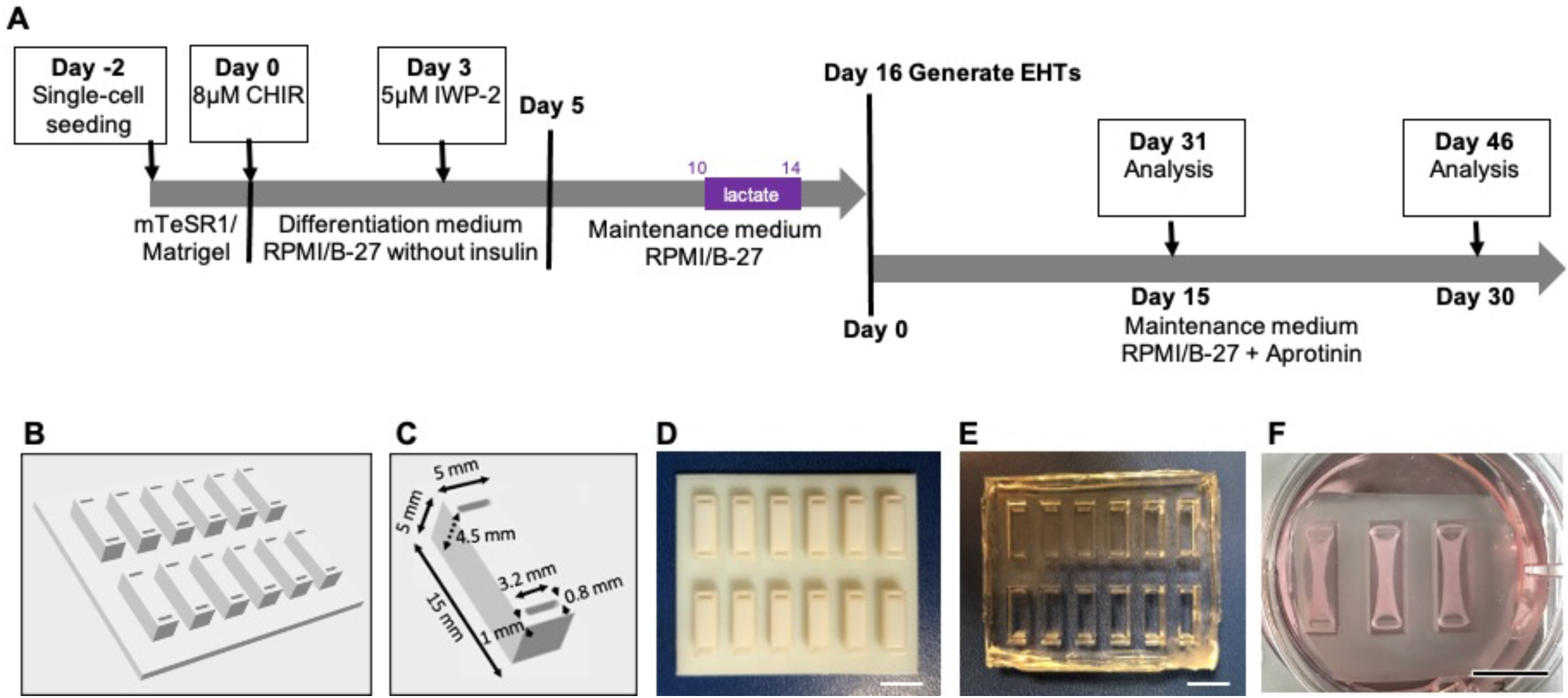
Fabrication of EHTs. (A) timeline of hiPSC-CM differentiation and EHT fabrication with key media composition labeled. (B and C) design and dimension of 3D-printed negative mold. (D and E) pictures of negative mold and casted PDMS mold. (Scale bar = 1 cm) (F) EHTs maintained in a PDMS mold inside a tissue culture well. (Scale bar = 1 cm)

EHTs were fabricated using a 3D printed negative mold (Figure 1B-D) into which polydimethylsiloxane (PDMS) was cast to form the EHT housing, termed “wells” (Figure 1E-F). Then a mixture comprising 1.6 million hiPSC-CMs and 0.4 million fibroblasts in fibrinogen solution was prepared, followed by the addition of thrombin before casting into PDMS wells. The cell solution immediately solidifies to form a fibrin-based structure that surrounds two posts. Within the initial 5 days following fabrication, the EHTs compact and undergo significant reduction in width, gradually reaching minimal width over 7 days. There were no discernible differences in EHT compaction between mutant and control groups. Consequently, parameters such as EHT length, width, and resulting passive strain remained consistent across both mutant and control EHTs (Supplementary Figure 1).

#### 1. 2. *MYH7/MYH6* mutant EHTs exhibit altered cardiomyocyte morphology, alignment, and calcium handling

To begin evaluating cardiomyocyte function within the EHT context, we used a calcium-sensitive dye to assess hiPSC-CM morphology and distribution. In both mutant and control EHTs, we observed predominant alignment of hiPSC-CMs along the long axis of the EHTs (see supplementary Figure 2A). However, while control EHTs exhibited well-integrated cardiomyocyte with contiguous muscle, mutant EHTs displayed isolated, discontinuous cardiomyocyte domains. This disparity became more pronounced by Day 30 post-EHT formation, with a normalized degree of alignment of 0.27 for Day 30 EHTs, as shown in supplementary Figure 2B. At Day 15, both mutant and control EHTs exhibited similar spontaneous beating rates (0.49Hz ± 0.021Hz and 0.43Hz ± 0.015Hz, respectively). However, by Day 30, the mutant EHTs maintained a consistent beat rate (0.42Hz ± 0.035Hz), while the control EHTs exhibited a significantly reduced spontaneous beat rate (0.27Hz ± 0.012Hz) as shown in Figure 2A. The declining trend in beat rate seen in the control EHTs suggests a progression from an embryo-like state to a more mature status during the development of immature hiPSC-CMs [35, 36]. However, a notable increase in irregular beating events was also observed in mutant EHTs (Figure 2B), consistent with our previous studies on HCM in cardiac monolayers [4]. Comparison of calcium signal amplitude revealed significantly increased amplitudes in mutant EHTs on both Day 15 and Day 30, suggesting that mutant cardiomyocytes recruited more calcium ions into the cells to complete each calcium handling cycle starting from Day 15. This increased calcium signal amplitude is also consistent with what we observed in 2D monolayer study [4]. We then evaluated the same calcium dynamic measures with 1Hz field pacing. We found that mutant and control EHTs displayed similar upstroke velocity at both time points and similar downstroke velocity at Day 15 (Figure 2D-2F). Over time, both mutant and control EHTs demonstrated progressive maturation, including increased upstroke and downstroke velocity. However, mutant EHTs exhibited significantly reduced downstroke velocity at Day 30 relative to controls.

**Figure 2.**
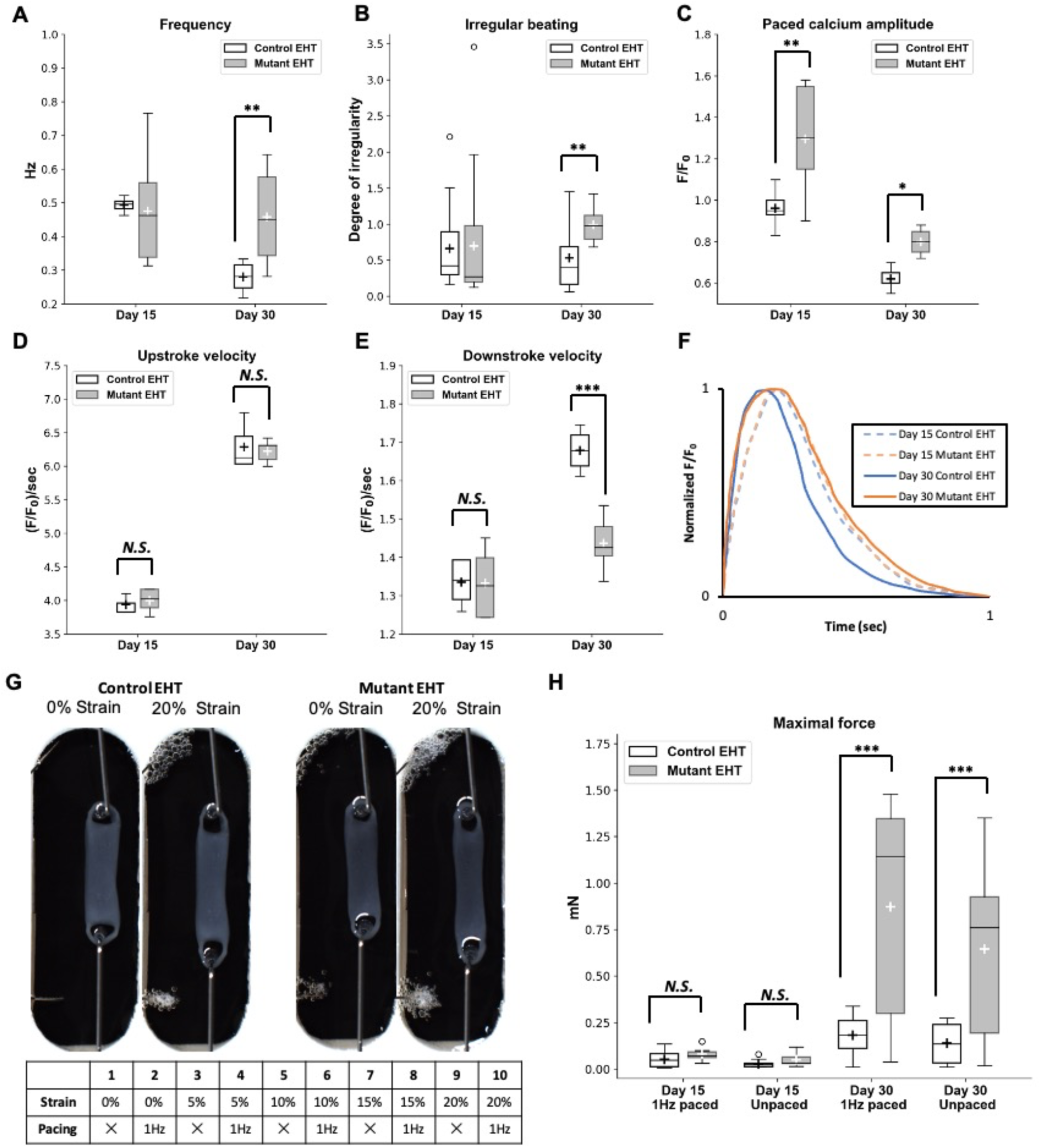
Functional alterations of mutant EHTs. (A) Beating rate measured from calcium transient analysis for mutant and control EHTs at Day 15 and 30 after fabrication. (*N.S.*: not significant; Student *t* test; three EHTs for each condition and two experimental replicates, n = 6 for both mutant and control groups) (B) Quantitative result for irregular beating of mutant and control EHTs at Day 15 and 30 after fabrication. (\*\**p* < 0.01; Student *t* test; three EHTs for each condition and two experimental replicates, n = 6 for both mutant and control groups) (C) Peak amplitude of calcium signal measured from calcium transient analysis with 1Hz field stimulation applied for mutant and control EHTs at Day 15 and 30 after fabrication. (\**p* < 0.05; \*\**p* < 0.01; Student *t* test; three EHTs for each condition and two experimental replicates, n = 6 for both mutant and control groups) (D) Upstroke velocity of calcium signal measured from calcium transient analysis with 1Hz field stimulation applied for mutant and control EHTs at Day 15 and 30 after fabrication. (*N.S.*: not significant; Student *t* test; three EHTs for each condition and two experimental replicates, n = 6 for both mutant and control groups) (E) Downstroke velocity of calcium signal measured from calcium transient analysis with 1Hz field stimulation applied for mutant and control EHTs at Day 15 and 30 after fabrication. (*N.S.*: not significant; \*\*\**p* < 0.005; Student *t* test; three EHTs for each condition and two experimental replicates, n = 6 for both mutant and control groups) (F) Representative calcium traces of calcium signal measured from calcium transient analysis with 1Hz field stimulation applied for mutant and control EHTs at Day 15 and 30 after fabrication. (G) Force generation measurements were assessed at Day 15 and 30 post-fabrication. During the force generation measurement, each EHT was incrementally stretched to 120% of its original length, and contraction forces were recorded under both 1Hz field-stimulated and unpaced conditions. (H) The maximum force of contraction of mutant and control EHTs (*N.S.*: not significant; \*\*\**p* < 0.005; Student *t* test; three EHTs for each condition and two experimental replicates, n = 6 for both mutant and control groups)

#### 1. 3. Force generation measurement reveals hypercontractility in mutant EHTs

One of the advantages of the EHT platform is the ability to directly assess cardiac mechanical function through force generation measurements. In this study, we compared mutant EHTs with control EHTs at two different time points, under both paced and unpaced conditions. Throughout these measurements, EHTs were incrementally stretched to 120% of the original length (20% strain, Figure 2G). Our findings highlighted that increased strain, achieved by stretching the EHTs, led to augmented contraction force in both mutant and control groups (supplementary Figure 3). Additionally, external electrical stimulation was applied to coordinate electromechanical function, enabling the investigation of the maximum summative force capacity of EHTs under different conditions (supplementary Figure 3). The result of pacing showed that both mutant and control EHTs adeptly matched a 1Hz pacing rate, with the peak force for all conditions manifesting under this 1Hz pacing.

A comparative analysis revealed that, irrespective of pacing and across both Day 15 and 30, mutant EHTs consistently demonstrated higher contraction force relative to the controls, consistent with the hypercontractility feature of HCM [37–40]. As shown in Figure 2H, the maximum force for Day 15 control EHTs was 0.049 ± 0.022mN, while Day 15 mutant EHTs registered at 0.081 ± 0.020mN. Similarly, the maximum force for Day 30 control EHTs was 0.182 ± 0.060mN, whereas the mutant EHTs reached a significantly higher force at 0.874 ± 0.330mN.

Consistent with observations from calcium transient measurements, the mutant EHTs exhibited notably faster spontaneous contraction and more irregular events. This phenomenon may be attributed to the presence of multiple beating sub-domains within the mutant EHTs, occasionally resulting in contractions offsetting each other. Consequently, when subjected to 1Hz pacing, the contraction force of the mutant EHTs not only stabilized but also showed significant augmentation under specific strain conditions. Furthermore, the discrepancy in contraction force between paced and unpaced conditions was more pronounced in mutant EHTs than the control EHTs, particularly by Day 30. This suggests that external electrical stimulation during force measurements assisted in coordinating electromechanical function further allowing the maximum summative force capacity of a given EHT. These results are distinct from our prior work with these same mutant cells in 2D [4], wherein mutant hiPSC-CMs showed a reduced contraction rate and compromised calcium handling.

Clinically, HCM is characterized by left ventricular (LV) hypertrophy, myocardial hypercontractility, diastolic dysfunction, reduced compliance, and either preserved or increased ejection fraction. At the cellular level, diseased cardiomyocytes exhibit hypertrophy, disorganization, and are interspersed with areas of interstitial fibrosis [41–43]. Our previous study of HCM using 2D cardiac monolayers revealed several cellular and molecular hallmarks of the disease and highlighted ECM dynamics as an early trigger of HCM. The 3D format, with the addition of fibroblasts, dramatically changed cardiomyocyte and resultant engineered tissue function, producing outcomes more consistent with the clinical manifestation of the disease. Through functional assessments, we delved into the cardiac functional changes caused by the MHC mutations. From here, we sought to unravel mechanistic determinants of disease onset and progression, especially in the context of cell-ECM engagement.

#### 1. 4. Mutant EHTs exhibit altered cell-ECM interaction and ECM composition

In our previous work, RNAseq analysis revealed pathways associated with ECM dynamics as the top discrepant pathways between mutant hiPSC-CMs and control hiPSC-CMs [4]. Here, we evaluated potential HCM mutation-related alterations in cell-ECM interactions in EHTs and started with IHC staining for pFAK (phosphorylated focal adhesion kinase) in hiPSC-CMs (Figure 3A). Results indicated a uniform distribution of hiPSC-CMs within the EHTs, and both mutant and control EHTs exhibited pFAK-positive signals. However, at Day 15, staining was more proximal to the edges in control EHTs, whereas mutant EHTs displayed more pronounced and dispersed pFAK signals across the EHT sections. Quantitative analysis of pFAK percent area revealed elevated levels of pFAK in mutant EHTs at Day 15, with no significant difference in expression between mutant and control EHTs at Day 30 (Figure 3B). Western blot analysis confirmed these findings, showing significantly elevated pFAK in mutant EHTs at Day 15, with no difference in expression between mutant and control EHTs at Day 30 (supplementary Figure 4A-4C). Notably, although the pFAK/FAK ratio showed an increasing trend from Day 15 to Day 30, the difference was not significant for both mutant and control EHTs. The difference between mutant and control at each time point was also not significantly different (Supplementary Figure 4D). Due to the nature of EHT, hiPSC-CMs and fibroblasts were mixed together. Therefore, IHC staining and Western blot analysis provided the pFAK protein levels expressed by both cell types without the ability to distinguish between cell types. To examine focal adhesions in cardiomyocytes exclusively, mutant and control hiPSC-CMs were cultured in a 2D format and transfected to express vinculin-GFP at 30 and 60 days after the initiation of CM differentiation [44, 45]. Vinculin is part of the focal adhesion complex, and was therefore used as a means to quantify focal adhesions in CMs. We found mutant hiPSC-CMs exhibited more focal adhesions per cell compared to controls. Although the difference was not statistically significant at Day 30 of differentiation (corresponding to Day 15 EHTs), it became significant at Day 60 of differentiation (Figure 3C and 3D). Together, these findings indicate alterations in ECM engagement with MHC mutations.

**Figure 3.**
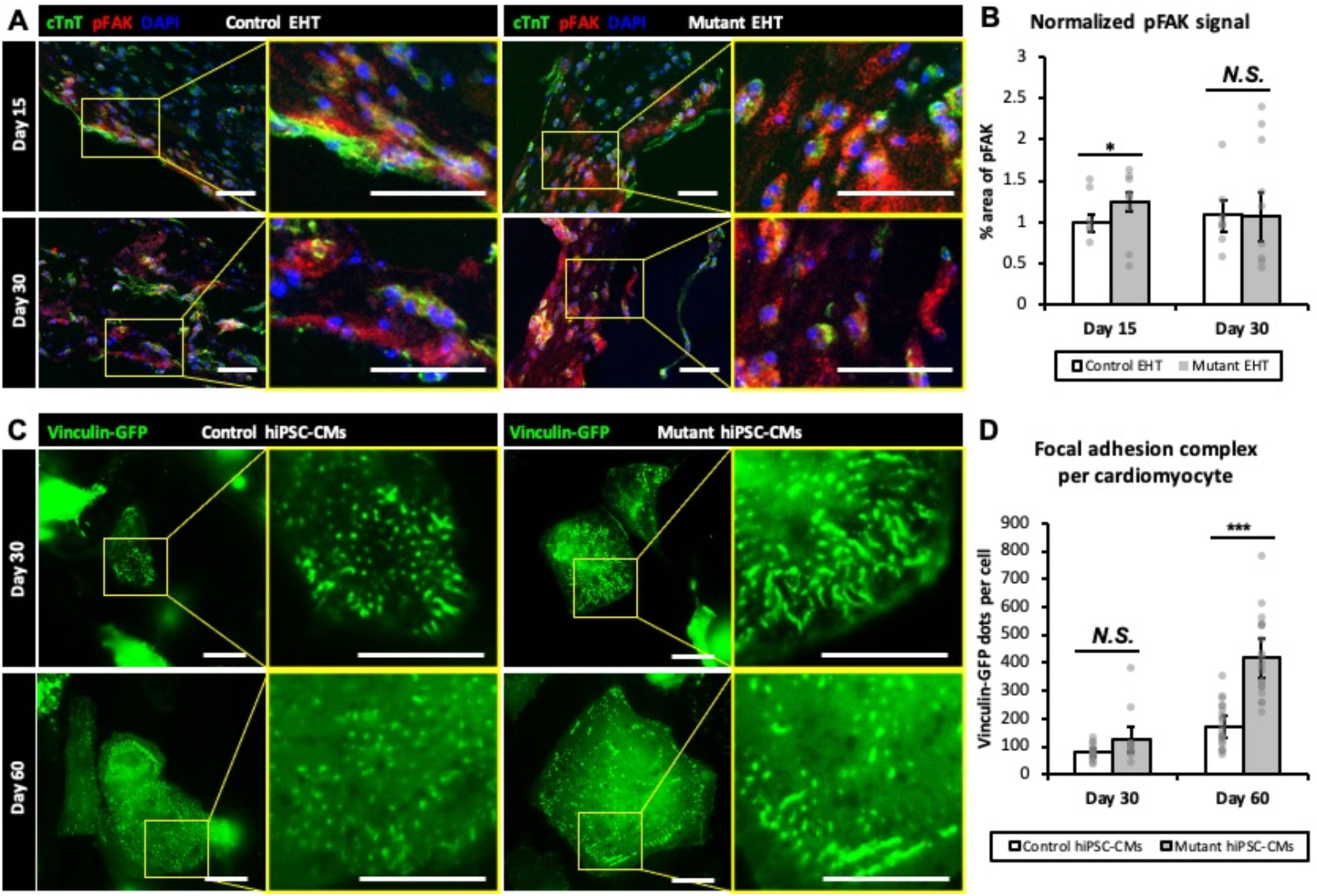
Altered cell-ECM engagement in mutant EHTs. (A) Representative images of cTnT, pFAK and DAPI-stained EHT sections for both control and mutant EHTs at Day 15 and 30. (Scale bar = 100 μm) (B) Quantitative analysis of pFAK percent area (normalized to Day 15 Control EHT). (*N.S.*: not significant; \**p* < 0.05; Student *t* test; n = 6 immunostained images for Day 15 control group, n = 9 for Day 30 control and mutant groups, and n = 8 for Day 15 mutant group) (C) Representative image of vinculin-GFP for control and mutant hiPSC-CMs at Day 30 and 60. (Scale bar = 25 μm) (D) Quantitative analysis of vinculin-GFP dots per cell for control and mutant hiPSC-CMs at Day 30 and Day 60. (\*\*\**p* < 0.005; *N.S.*: not significant; Student *t* test; n = 15 cells for Day 30 control condition, n = 12 cells for Day 30 mutant condition, n = 22 cells for Day 60 control condition, and n = 20 cells for Day 60 mutant condition)

To determine whether changes in focal adhesions were associated with the same or different type of ECM in the EHT, we evaluated the composition of ECM in EHTs using Masson’s trichrome and Alcian blue staining. As shown in Figure 4A, Masson’s trichrome staining indicated that both control and mutant EHTs began producing collagen by Day 30, with no significant difference in collagen levels between the two genotypic conditions (Figure 4A and 4B). Notably, muscle tissues appeared as a darker red color, with control EHTs showing organized and connected muscle tissue by Day 15, whereas mutant EHTs exhibited a sparse pattern of muscle tissue. Interestingly, Alcian blue staining revealed a significantly higher presence of ECM proteoglycans in control EHTs at Day 30 relative to mutant cells (Figure 4A and 4C). Given this difference, we probed a bit more deeply using IHC for key fibrosis-associated ECM, type I collagen and fibronectin (Figure 4D). The quantitative analysis of IHC images indicated no discernible difference between the control and mutant groups in terms of type I collagen and fibronectin levels on both Day 15 and 30, consistent with the colorimetric stains (Figure 4E and 4F). These results suggest ECM remodeling at the early stage precedes and is distinct from late-stage fibrosis associated with HCM.

**Figure 4.**
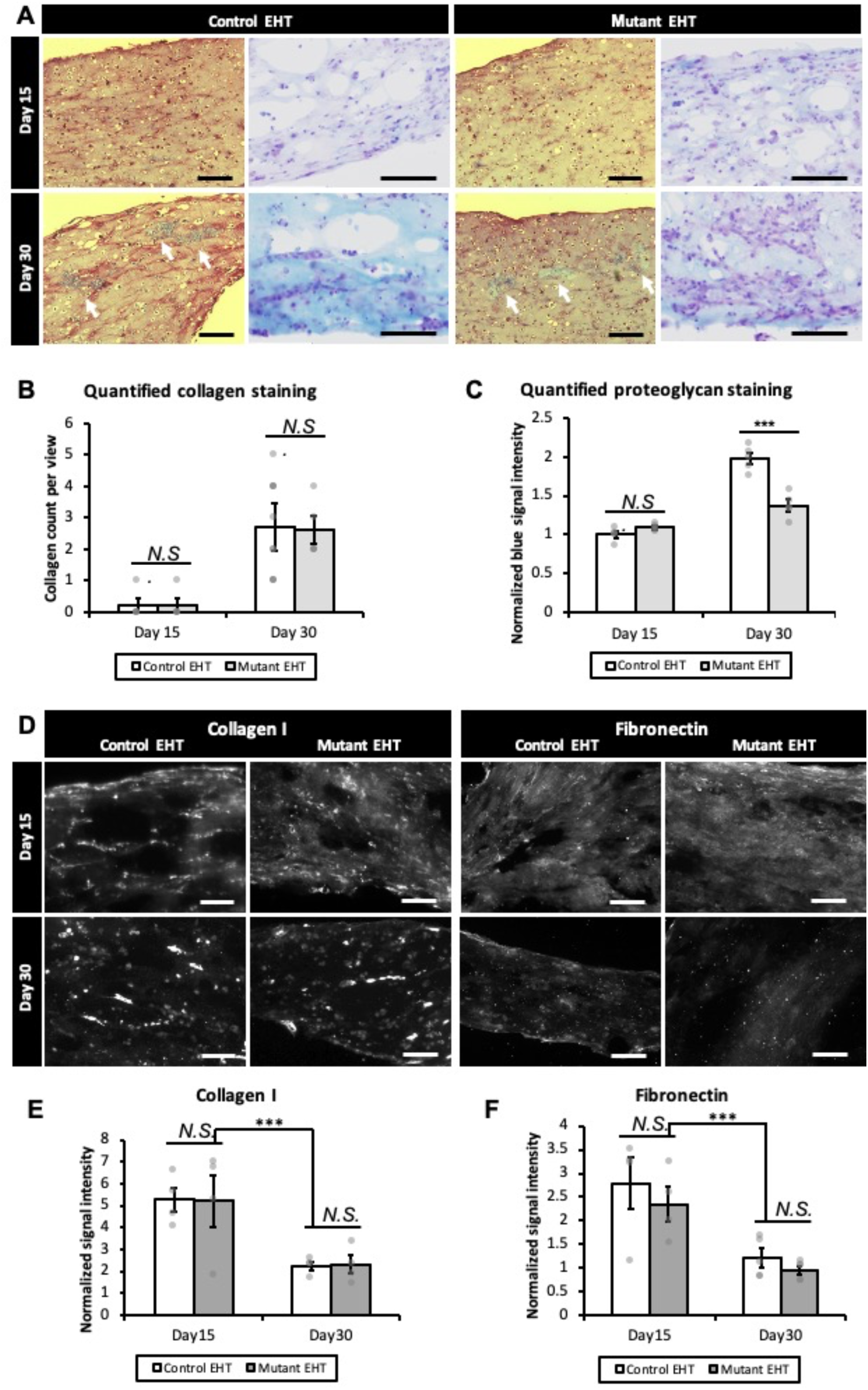
Altered ECM composition in mutant EHTs. (A) Masson’s trichrome (right panel) and Alcian blue (left panel) staining of EHT sections from both control and mutant samples at Day 15 and 30. White arrows highlight areas of deposited collagen. (Scale bar = 100 μm) (B) Quantitative analysis of deposited collagen per section area. (*N.S.*: not significant; Student *t* test; n = 6 immunostained images for each condition) (C) Quantitative analysis of proteoglycan signal intensity per section area. (*N.S.*: not significant; \*\*\**p* < 0.005; Student *t* test; n = 6 immunostained images for each condition) (D) Representative image of maximum intensity of collagen I and Fibronectin stained EHT sections from both control and mutant samples at Day 15 and 30. (Scale bar = 100 μm) (E) Quantitative analysis of collagen I signal intensity per section area. (*N.S.*: not significant; Student *t* test; n = 5 immunostained images for each condition) (F) Quantitative analysis of fibronectin signal intensity per section area. (*N.S.*: not significant; Student *t* test; n = 5 immunostained images for each condition)

#### 1. 5. Increased percentage of activated fibroblasts and elevated TGF-β1 secretion in the EHTs with mutant hiPSC-CM

To determine whether altered activation of fibroblasts could account for differential expression of ECM in the mutant case, we stained for αSMA+/cTnT-activated fibroblasts (Figure 5A). Activated fibroblasts primarily congregated near the edges of control EHTs, whereas mutant EHTs displayed a more uniform distribution of activated fibroblasts, consistent with the pattern observed for pFAK signals. Quantitative analysis of immunostaining images showed an increased population of activated fibroblasts in mutant EHTs as early as Day 15. Control EHTs exhibited 43.78% ± 3.10% and 46.73% ± 4.63% of activated fibroblast at Day 15 and 30 respectively, whereas mutant EHTs displayed 56.78% ± 5.91% and 54.99% ± 2.35% of activated fibroblast at Day 15 and 30 respectively (Figure 5B).

**Figure 5.**
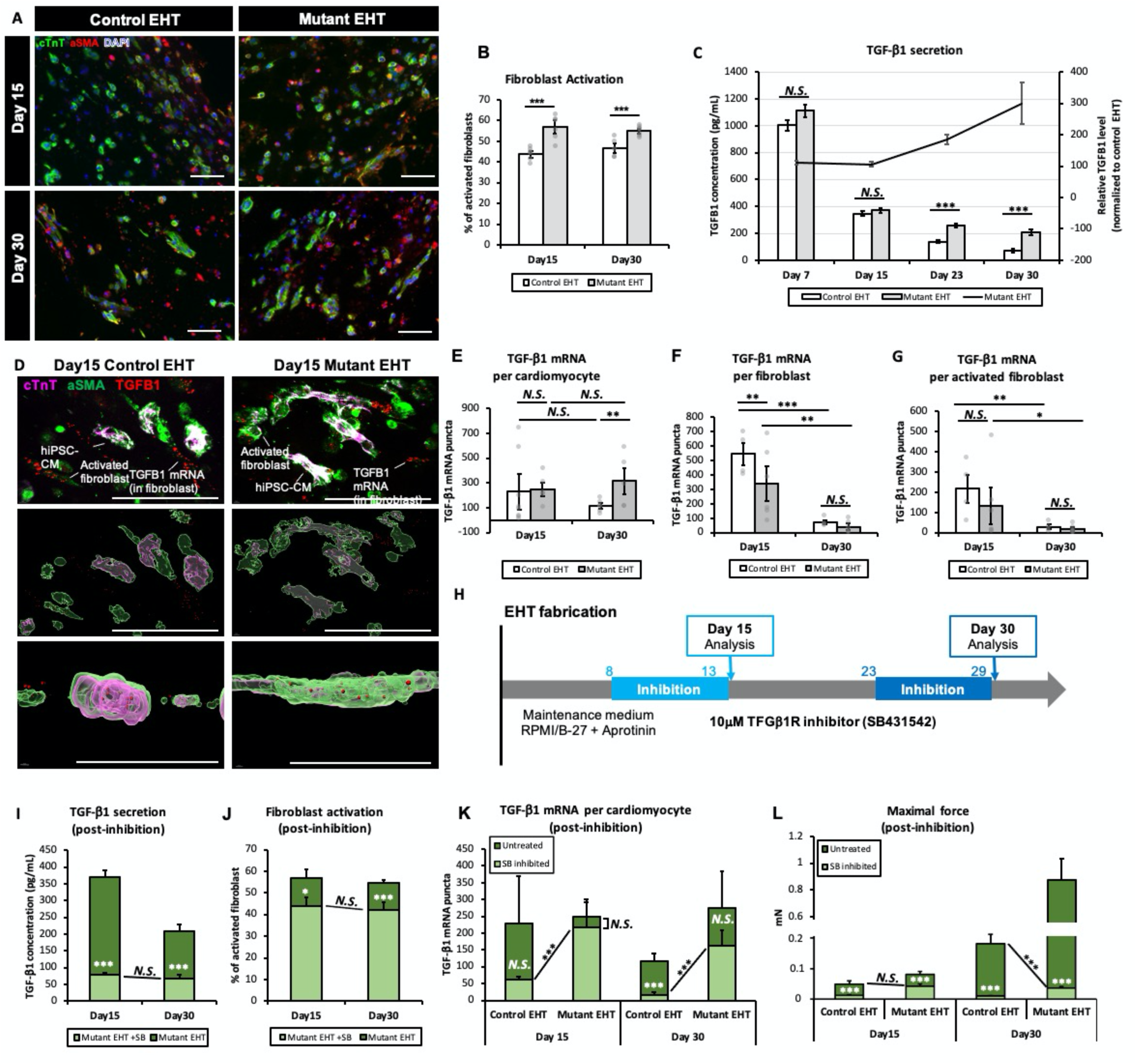
Pathological upregulation of TGF-β1 expression by mutant hiPSC-CMs and the efficacy of TGF-β1R inhibition in attenuating HCM-associated hypercontractility. (A) Representative images of cTnT, aSMA and DAPI-stained EHT sections from both control and mutant samples at Day 15 and 30. (Scale bar = 100 μm) (B) Quantitative analysis of activated fibroblasts relative to the total fibroblast population in both control and mutant EHTs at Day 15 and 30. (\*\*\**p* < 0.005; Student *t* test; n = 6 immunostained images for each condition) (C) TFG-β1 secretion via ELISA analysis of exhausted medium collected from control and mutant EHT wells at Day 7, 15, 23 and 30. (*N.S.*: not significant; \*\*\**p* < 0.005; Student *t* test; three wells for each condition and 2 experimental replicates, n = 6 for both control and mutant groups) (D) Representative images of cTnT, αSMA, and TGF-β1-stained sections from both control and mutant EHT samples at Day 15 and 30 (top panel). Imaris software captured cTnT, αSMA, and TGF-β1 signals (middle panel), along with z-axis images (bottom panel) demonstrating the presence of TGF-β1 puncta within cells. (Scale bar = 100 μm) (E) Quantitative analysis of TGF-β1 mRNA expression per cardiomyocyte in both control and mutant EHTs at Day 15 and 30. (\*\**p* < 0.01; Student *t* test; n = 6 FISH-IHC stained images for each condition) (F) Quantitative analysis of TGF-β1 mRNA expression per fibroblast in both control and mutant EHTs at Day 15 and 30. (\*\**p* < 0.01; \*\*\**p* < 0.005; Student *t* test; n = 6 FISH-IHC stained images for each condition) (G) Quantitative analysis of TGF-β1 mRNA expression per activated fibroblast in both control and mutant EHTs at Day 15 and 30. (\**p* < 0.05; \*\**p* < 0.01; Student *t* test; n = 6 FISH-IHC stained images for each condition) (H) TGF-β1R inhibition timeline and regime. (I) Post-inhibition TGF-β1 secretion comparing to untreated levels in exhausted medium collected from mutant EHT wells at Day 15 and 30. (*N.S.*: not significant; \*\*\**p* < 0.005; Student *t* test; three wells for each condition and 2 experimental replicates, n = 6 for both control and mutant groups) (J) Post-inhibition percentage of activated fibroblasts comparing to untreated level in both control and mutant EHTs at Day 15 and 30. (\**p* < 0.05; \*\*\**p* < 0.005; *N.S.*: not significant; Student *t* test; n = 6 immunostained images for each condition) (K) Post-inhibition TGF-β1 mRNA expression per cardiomyocyte in both control and mutant EHTs at Day 15 and 30. (\*\*\**p* < 0.005; *N.S.*: not significant; Student *t* test; n = 6 FISH-IHC stained images for each condition) (L) Post-inhibition maximum force of contraction comparing to untreated level in mutant and control EHTs at Day 15 and 30. (*N.S.*: not significant; \*\*\**p* < 0.005; Student *t* test; three EHTs for each condition and two experimental replicates, n = 6 for both mutant and control groups)

There are several pathways that govern fibroblast activation, and TGF-β1-driven fibroblast activation is reported to be one essential pathway in cardiac tissues in later stage disease [42, 43, 46–48]. Given the observed upsurge in contraction force and the increased presence of activated fibroblasts in mutant EHTs, we conducted a TGF-β1 ELISA to determine if cell-cell interactions might be mediated through TGF-β1 signaling in this HCM model. We collected exhausted media from both mutant and control EHTs at various time points and analyzed the TGF-β1 concentration (Figure 5C). The result showed that TGF-β1 concentration decreased over time for both mutant and control EHTs. However, the mutant EHTs had a slower rate of decrease of TGF-β1 concentration over time. Consequently, from Day 23 of EHT fabrication onward, TGF-β1 secretion was notably elevated in mutant EHTs compared to the controls.

To discern which cell type was responsible for TGF-β1 secretion, we employed RNA fluorescent in situ hybridization (RNA FISH) using a commercial TGFβ1 probe, adhering to the RNAscope protocol. In both mutant and control EHTs, a punctate pattern of TGF-β1 mRNA signals was observed within the cytosol (Figure 5D). We also found significantly more TGF-β1 mRNA molecules in mutant cardiomyocytes on Day 30 when compared to the control (Figure 5E). Despite fibroblasts exhibiting a higher initial number of TGF-β1 mRNA molecules per cell than hiPSC-CMs, there was surprisingly no discernible difference in the TGF-β1 mRNA molecules generated by either fibroblasts or activated fibroblasts between the control and mutant EHTs at the later time point (Figure 5F). Consequently, the heightened TGF-β1 mRNA levels in mutant EHTs at Day 30 could primarily be attributed to the mutant hiPSC-CMs.

#### 1. 6. TGF-β1 receptor inhibition mitigates increased TGF-β1 secretion and force of contraction in mutant EHTs through disruption of intercellular communication between mutant hiPSC-CMs and fibroblasts

To examine whether blocking TGF-β1 signaling could attenuate the pathological changes in ECM secretion and force of contraction of mutant EHTs, we employed SB431542, a small molecule inhibitor of the TGF-β1 receptor, to block the interplay between cardiomyocyte and fibroblasts. As shown in Figure 5H, a concentration of 10 µM of SB431542 was introduced to the culture medium for a duration of 7 days preceding the designated analysis time point (e.g., from Day 8 to 15 for Day 15 analysis and from Day 23 to 30 for Day 30 analysis). TGF-β1 secretion in the absence of the inhibitor showed no difference between mutant and control EHTs at Day 15 (Figure 5C). However, post-inhibition, TGF-β1 secretion in Day 15 mutant EHTs plummeted to 21% relative to its untreated counterparts. By Day 30, untreated mutant EHTs exhibited markedly higher TGF-β1 secretion compared to the controls. Yet, upon inhibition, the secretion in mutant EHTs dropped further, aligning closely with the secretion levels observed in Day 30 control EHTs (Figure 5I). Further IHC staining of αSMA, cTnT, and DAPI revealed that both Day 15 and Day 30 mutant EHTs plus TGFβ1R inhibitor possessed fewer αSMA+/cTnT-(i.e., activated fibroblasts) than their untreated counterparts at the respective time points, suggesting a decrease in the number of activated fibroblasts post TGF-β1R inhibition (Figure 5J). Interestingly, on a per hiPSC-CM basis, the number of TGF-β1 mRNA molecules remained consistently elevated in the mutant EHTs. This finding indicated that blockage of TGF-β1 receptor inhibits paracrine signals, despite the fact that mutant hiPSC-CMs were still generating higher amounts of TGF-β1 mRNA (Figure 5K). In terms of contraction force, mutant EHTs subjected to inhibition displayed a significant decrease. By Day 15, the inhibited mutant EHTs exhibited contraction forces on par with Day 15 control EHTs. By Day 30, these forces in the inhibited mutant EHTs diminished to roughly one-fifth of those observed in the untreated Day 30 control EHTs (Figure 5H). The decrease in contraction force following TGF-β1R inhibition was not due to cell death or loss of contractile capacity. As shown in supplementary Figure 5, although the contraction force significantly decreased for the Day 30 mutant EHTs, there was no discernible difference in the IHC images regarding total cell number and morphology between the untreated and TGF-β1R inhibited groups.

## Discussion

Using a multicellular engineered heart tissue model, we find that the *early* interplay of MHC mutant hiPSC-CMs with fibroblasts drive dysfunctional features associated with HCM and that this interplay is governed in part by TGF-β1 signaling by *cardiomyocytes* with respondent fibroblast activation. These mechanistic findings provide strong scientific premise for developing and implementing novel strategies to treat HCM well before the manifestation of clinically detectable cardiac dysfunction or fibrosis.

Much of what we know about HCM-related ECM changes corresponds to fibrosis at late stages of disease progression when the disease is detected clinically via tissue scale imaging [49–53]. However, recently we used hiPSC-CM to detect altered ECM dynamics very soon after the specification of hiPSC-CMs carrying HCM-associated genetic mutations [4]. Here, we aimed to delve into the underlying mechanisms governing cell-ECM engagement in early-stage HCM using a multicellular (i.e., CM and fibroblasts) EHT model. To examine the early-stage HCM-associated ECM changes in our EHT system, we conducted histological staining for ECM proteins and immunostaining for cellular components reflective of cell-ECM engagement. We found mutant EHTs exhibited significantly lower levels of proteoglycans by Day 30. Proteoglycans have been reported to promote favorable cardiac remodeling after pressure overload and mitigate pathological conditions [55]. Additionally, the accumulation of proteoglycans is associated with post-inflammatory remodeling [55]. The increased abundance of proteoglycans in control EHTs may signify a compensatory mechanism, wherein the accumulated proteoglycans mitigate increased physiological cues (such as passive strain resulting from the EHT model design) and assist in healthy cardiac tissue development. Mutant EHTs did not show proteoglycan accumulation, suggesting they followed a pathological route. This outcome is consistent with our 2D study showing a decrease in RNA transcript for all ECM genes differentially expressed between mutant and control hiPSC-CM [4] and a study by Larson et al., showing reduced ECM gene expression in cardiomyocytes of HCM patients [50]. Concordant with altered ECM expression, mutant EHTs exhibited an altered means of engaging with the varied ECM of the mutant as evidenced by more intense and evenly distributed pFAK signals throughout the tissue, relative to controls. Further, mutant hiPSC-CMs had more focal adhesions per cell than controls. We hypothesized that differential ECM expression and associated ECM engagement arose as a function of altered fibroblast activation and associated ECM remodeling.

Previous studies have highlighted the crucial role of fibroblast activation in heart development and various cardiac diseases, including HCM [29, 56–59]. Furthermore, evidence suggests that activation of fibrotic pathways precedes the onset of hypertrophic remodeling, indicating that fibroblast activation is an upstream factor in the etiology of HCM [39, 59, 60]. Indeed, we observed an increased relative fraction of activated fibroblasts in mutant EHTs at early time points, potentially associated with the initiation of disease development. Further, it has been reported that isoforms of TGF-β trigger the essential signaling pathways of fibroblast activation and therefore play a central role in the onset of fibrosis. [42, 43, 48, 59, 61]. Multiple types of cells of cardiac tissue, such as CMs, fibroblasts, endothelial cells and immune cells, can secrete TGF-β [59, 61, 62]. Once TGF-β binds to surface receptors on the fibroblasts, intracellular signaling cascades will be initiated and further trigger the expression of α-SMA, regulate the expression of genes involved in ECM turnover, and the consequent interstitial fibrosis [59, 63–65]. Through plasma proteomic profiling and analysis, significantly increased serum TGF-β1 levels in HCM patients have been found. Here we demonstrated through ELISA assessment that media from the mutant culture wells had significantly higher TGF-β1 concentrations than those from the control wells, starting as early as Day 23 of CM differentiation. Moreover, the sustained level of TGF-β1 secretion in mutant EHTs suggest that the interaction between mutant hiPSC-CMs and fibroblasts likely operates, at least in part, through TGF-β1 signaling. Many studies indicate an increased gene or protein expression of TGF-β1 in HCM patients and the correlation to disease onset and severe outcome [58, 65–68]; however, the mechanism *driving TGF-β1 expression with HCM* remains unknown.

In this study, we utilized RNA FISH to investigate TGF-β1 expression in fibroblasts and cardiomyocytes in this early-stage HCM model. Interestingly, the number of TGF-β1 mRNA transcripts within fibroblasts was not affected by the different genetic mutations of the neighboring hiPSC-CMs at both time points (supplementary Figure 6). Similarly, the number of TGF-β1 mRNA transcripts per cell showed no difference between mutant and control hiPSC-CMs on Day 15. However, by Day 30, despite the overall drop in TGF-β1 expression levels, mutant hiPSC-CMs retained expression of TGFβ1 mRNA. This likely contributed to the significantly elevated TGF-β1 protein secretion observed in mutant EHTs. Teekakirikul et al. employed mouse models carrying human HCM mutations and showed enhanced TGF-β1 expression in non-cardiomyocytes [64]. The differences between their findings and ours could be primarily due to the modeling timescale, as our EHT model was designed to recapitulate early-stage HCM well before the onset of clinical symptoms. Altogether, cardiomyocytes utilized TGF-β1 signals to activate and interface with supporting cells in the early HCM pathology. The findings from the RNA FISH analysis aligned with the ELISA, activated fibroblasts, and force generation results. Specifically, MHC mutant CM not only produced an increased amount of TGF-β1 mRNA but also secreted a higher quantity of TGF-β1 proteins into the culture medium.

When considering the factors that might drive TGF-β expression in cardiomyocytes, we start with the impact of the mutation itself on cell function. It is known that sarcomere protein mutations can result in enhanced biomechanical forces, which could turn on TGF-β signaling and further activate neighboring non-cardiomyocytes [65–67]. Moreover, biophysical studies have shown that MHC mutations cause aberrant slipping of the myosin head past the actin filament more rapidly than the wild type case [68]. More rapid contraction could generate more force or tension on the costamere and associated focal adhesion of the cardiomyocytes. From here, there is evidence to suggest that activation of ERK, downstream of FAK activation, can permit ERK to transit to the nucleus and drive the expression of TGF-β. To determine whether blocking TGF-β1 signaling limits disease progression, we conducted chronic TGF-β1R inhibition. Indeed, inhibition of the receptor decreased the contraction force, TGF-β1 concentration in culture medium, and the percentage of activated fibroblasts in mutant EHTs to levels consistent with control EHTs. Blocking TGF-β1 signaling by masking the receptor was sufficient to rescue the mutant EHTs, indicating the potential therapeutic efficacy of targeting this pathway in HCM. To further determine whether aberrant biophysical contraction caused by the mutation drives altered CM-ECM engagement and corresponding changes in MAPK/ERK signaling, one could introduce an inhibitor of FAK phosphorylation and evaluate the consequent activation of ERK with MHC mutation.

The clinical management of HCM patients presents challenges due to the heterogeneous clinical expression and mutation-specific pathological features associated with the disease. This is exacerbated by the fact that we understand very little about early-stage pathology. Current pharmacological therapy for HCM patients includes β-adrenergic receptor blockers (β-blockers), non-dihydropyridine calcium channel blockers (CCBs), and disopyramide [73, 74]. β-blockers, the first-line therapy of HCM for decades, mitigate exercise-induced left ventricular outflow tract obstruction (LVOTO) and relieve symptoms like arrhythmias and myocardial hypercontractility. However, β-blockers do not improve prognosis or reduce sudden death rates in HCM patients [75]. Non-dihydropyridine CCBs, which block the L-type calcium channel and reduce calcium overload, are effective when β-blockers are poorly tolerated. Disopyramide reduces early flow velocity in the LVOT and mitigates systolic anterior motion of the anterior mitral leaflet, making it effective in reducing heart failure symptoms and LVOT gradient [76, 77]. However, these non-selective drugs primarily target late disease manifestations without addressing intrinsic pathological mechanisms, rendering them effective for only a minority of patients [73, 78]. Given that HCM patients are reported to exhibit hypercontractility and impaired relaxation features [79–81], novel compounds that directly inhibit myosin have been developed. [73, 78]. For instance, Mavacamten administration decreases ATPase activity, thereby reducing the power output generated by the sarcomere [82, 83]. However, the preventive effect may be limited to certain mutations and dependent on the patient’s energy and activity levels [84].

Building on these pharmacological approaches, our study proposes targeting cardiomyocyte-induced fibroblast activation in the early stages of HCM to delay disease onset. As described above, TGF-β1 signaling is initiated as a result of sarcomere mutation, enhanced biomechanical forces, and the activation of FAK and ERK, providing several therapeutic targets for intervening HCM pathology. Animal studies and clinical trials have tested inhibitors targeting TGF-β, FAK or ERK pathways to prevent fibrosis progression. Angiotensin receptor blockers can inhibit TGF-β, and a phase II trial showed its potential in improving cardiac structure and function at early stages of HCM [85–87]. PF-573228 is a potent and selective FAK inhibitor that significantly attenuated fibrosis and preserved heart function *in vivo* [88]. MEK inhibitor, Mirdametinib, which blocks the ERK pathway, normalized the HCM-associated cardiac defects in a transgenic mouse model [89, 90]. However, it is challenging to achieve a balance between efficacy and off-target toxicity in non-cardiac cells when blocking these pathways in a non-cell type-specific manner. Further, the effectiveness of this strategy is limited to the early stage of the disease and may not effectively intervene in patients where fibrosis and pathological alterations have already commenced. Therefore, we propose targeting cardiomyocyte-specific pathological expression of TGF-β1 at the early-stage of HCM progression through RNA-based therapeutics to directly lower TGF-β1 expression and disrupt the downstream pathways.

In conclusion, our findings highlight the role of TGF-β1 signaling via MHC mutant cardiomyocytes in mediating the early onset of HCM. Blocking TGF-beta-mediated signaling may reverse early-stage functional alterations and extracellular matrix changes associated with HCM. Future therapeutic approaches may involve RNA-based therapy combined with targeted drug delivery to directly target pathways contributing to elevated TGF-β1 mRNA in mutant cardiomyocytes, offering promising prospects for HCM treatment.

## Acknowledgements

This research was supported by the American Heart Association (#21Predoc836194 to JH) and Regenerative Medicine Minnesota (RMM 091620DS008 to BO).

## Author contributions

JH and BMO conceived and designed the research. JH designed and executed the experiments, including cardiac differentiation, EHT fabrication, functional measurements, TGFB1R inhibition experiments, endpoint sample collection and slide perpetration, immunostaining, imaging and quantitative analysis, as well as ELISA experiments and analysis. JH, MALH and RLM designed the RNA FISH experiments. JH and MALH conducted the RNA FISH experiments and analysis. MS performed the Western blot experiments. FK and PJE design the vinculin-GFP experiment. MS performed cardiac differentiation and seeded hiPSC-CMs on Matrigel, and PJE performed vinculin-GFP transfection and imaging. MS produced the quantitative result of vinculin-GFP. JH and BMO discussed the results, wrote, and edited the manuscript. All authors approved the manuscript.

## Competing interests

The authors declare no competing interests.

## Notes

### Competing Interest Statement

The authors have declared no competing interest.

## References

1. Semsarian C, Ingles J, Maron MS, Maron BJ. New Perspectives on the Prevalence of Hypertrophic Cardiomyopathy, Journal of the American College of Cardiology, Volume 65, Issue 12, 2015, Pages 1249–1254, ISSN 0735-1097, doi: 10.1016/j.jacc.2015.01.019.

2. Maron BJ, Gardin JM, Flack JM, Gidding SS, Kurosaki TT, Bild DE. Prevalence of hypertrophic cardiomyopathy in a general population of young adults. Echocardiographic analysis of 4111 subjects in the CARDIA Study. Coronary Artery Risk Development in (Young) Adults. Circulation. 1995 Aug 15;92(4):785–9. doi: 10.1161/01.cir.92.4.785. PMID: 7641357.

3. Wolf CM. Hypertrophic cardiomyopathy: genetics and clinical perspectives. Cardiovasc Diagn Ther. 2019 Oct;9(Suppl 2):S388-S415. doi: 10.21037/cdt.2019.02.01. PMID: 31737545; PMCID: PMC6837941.

4. Hsieh J, Becklin KL, Givens S, Komosa ER, Lloréns JEA, Kamdar F, Moriarity BS, Webber BR, Singh BN, Ogle BM. Myosin Heavy Chain Converter Domain Mutations Drive Early-Stage Changes in Extracellular Matrix Dynamics in Hypertrophic Cardiomyopathy. Front Cell Dev Biol. 2022 Jun 16;10:894635. doi: 10.3389/fcell.2022.894635. PMID: 35784482; PMCID: PMC9245526.

5. Andrysiak K, Machaj G, Priesmann D, Woźnicka O, Martyniak A, Ylla G, Krüger M, Pyza E, Potulska-Chromik A, Kostera-Pruszczyk A, Łoboda A, Stępniewski J, Dulak J. Dysregulated iron homeostasis in dystrophin-deficient cardiomyocytes: correction by gene editing and pharmacological treatment. Cardiovasc Res. 2024 Feb 27;120(1):69–81. doi: 10.1093/cvr/cvad182. PMID: 38078368; PMCID: PMC10898935.

6. Li, J., Feng, X. & Wei, X. Modeling hypertrophic cardiomyopathy with human cardiomyocytes derived from induced pluripotent stem cells. Stem Cell Res Ther 13, 232 (2022). doi: 10.1186/s13287-022-02905-0.

7. Nguyen AH, Marsh P, Schmiess-Heine L, Burke PJ, Lee A, Lee J, Cao H. Cardiac tissue engineering: state-of-the-art methods and outlook. J Biol Eng. 2019 Jun 28;13:57. doi: 10.1186/s13036-019-0185-0. PMID: 31297148; PMCID: PMC6599291.

8. Tani H, Tohyama S. Human Engineered Heart Tissue Models for Disease Modeling and Drug Discovery. Front Cell Dev Biol. 2022 Mar 31;10:855763. doi: 10.3389/fcell.2022.855763. PMID: 35433691; PMCID: PMC9008275.

9. Cashman TJ, Josowitz R, Johnson BV, Gelb BD, Costa KD. Human Engineered Cardiac Tissues Created Using Induced Pluripotent Stem Cells Reveal Functional Characteristics of BRAF-Mediated Hypertrophic Cardiomyopathy. PLoS One. 2016 Jan 19;11(1):e0146697. doi: 10.1371/journal.pone.0146697. PMID: 26784941; PMCID: PMC4718533.

10. Wang K, Schriver BJ, Aschar-Sobbi R, Yi AY, Feric NT, Graziano MP. Human engineered cardiac tissue model of hypertrophic cardiomyopathy recapitulates key hallmarks of the disease and the effect of chronic mavacamten treatment. Front Bioeng Biotechnol. 2023 Sep 8;11:1227184. doi: 10.3389/fbioe.2023.1227184. PMID: 37771571; PMCID: PMC10523579.

11. Burke MA, Cook SA, Seidman JG, Seidman CE. Clinical and Mechanistic Insights Into the Genetics of Cardiomyopathy. J Am Coll Cardiol. 2016 Dec 27;68(25):2871–2886. doi: 10.1016/j.jacc.2016.08.079. PMID: 28007147; PMCID: PMC5843375.

12. Glazier AA, Thompson A, Day SM. Allelic imbalance and haploinsufficiency in MYBPC3-linked hypertrophic cardiomyopathy. Pflugers Arch. 2019 May;471(5):781–793. doi: 10.1007/s00424-018-2226-9. Epub 2018 Nov 20. PMID: 30456444; PMCID: PMC6476680.

13. Montag J, Syring M, Rose J, Weber AL, Ernstberger P, Mayer AK, Becker E, Keyser B, Dos Remedios C, Perrot A, van der Velden J, Francino A, Navarro-Lopez F, Ho CY, Brenner B, Kraft T. Intrinsic MYH7 expression regulation contributes to tissue level allelic imbalance in hypertrophic cardiomyopathy. J Muscle Res Cell Motil. 2017 Aug;38(3-4):291–302. doi: 10.1007/s10974-017-9486-4. Epub 2017 Nov 3. PMID: 29101517; PMCID: PMC5742120.

14. Burkart V, Kowalski K, Aldag-Niebling D, Beck J, Frick DA, Holler T, Radocaj A, Piep B, Zeug A, Hilfiker-Kleiner D, Dos Remedios CG, van der Velden J, Montag J, Kraft T. Transcriptional bursts and heterogeneity among cardiomyocytes in hypertrophic cardiomyopathy. Front Cardiovasc Med. 2022 Aug 23;9:987889. doi: 10.3389/fcvm.2022.987889. PMID: 36082122; PMCID: PMC9445301.

15. Litviňuková M, Talavera-López C, Maatz H, Reichart D, Worth CL, Lindberg EL, Kanda M, Polanski K, Heinig M, Lee M, Nadelmann ER, Roberts K, Tuck L, Fasouli ES, DeLaughter DM, McDonough B, Wakimoto H, Gorham JM, Samari S, Mahbubani KT, Saeb-Parsy K, Patone G, Boyle JJ, Zhang H, Zhang H, Viveiros A, Oudit GY, Bayraktar OA, Seidman JG, Seidman CE, Noseda M, Hubner N, Teichmann SA. Cells of the adult human heart. Nature. 2020 Dec;588(7838):466-472. doi: 10.1038/s41586-020-2797-4. Epub 2020 Sep 24. PMID: 32971526; PMCID: PMC7681775.

16. Grandi E, Navedo MF, Saucerman JJ, Bers DM, Chiamvimonvat N, Dixon RE, Dobrev D, Gomez AM, Harraz OF, Hegyi B, Jones DK, Krogh-Madsen T, Murfee WL, Nystoriak MA, Posnack NG, Ripplinger CM, Veeraraghavan R, Weinberg S. Diversity of cells and signals in the cardiovascular system. J Physiol. 2023 Jul;601(13):2547–2592. doi: 10.1113/JP284011. Epub 2023 Feb 16. PMID: 36744541; PMCID: PMC10313794.

17. Owen TJ, Harding SE. Multi-cellularity in cardiac tissue engineering, how close are we to native heart tissue? J Muscle Res Cell Motil. 2019 Jun;40(2):151–157. doi: 10.1007/s10974-019-09528-8. Epub 2019 Jun 20. PMID: 31222588; PMCID: PMC6726707.

18. Mourad O, Yee R, Li M, Nunes SS. Modeling Heart Diseases on a Chip: Advantages and Future Opportunities. Circ Res. 2023 Feb 17;132(4):483–497. doi: 10.1161/CIRCRESAHA.122.321670. Epub 2023 Feb 16. PMID: 36795846.

19. Munawar S, Turnbull IC. Cardiac Tissue Engineering: Inclusion of Non-cardiomyocytes for Enhanced Features. Front Cell Dev Biol. 2021 May 25;9:653127. doi: 10.3389/fcell.2021.653127. PMID: 34113613; PMCID: PMC8186263.

20. Voges HK, Foster SR, Reynolds L, Parker BL, Devilée L, Quaife-Ryan GA, Fortuna PRJ, Mathieson E, Fitzsimmons R, Lor M, Batho C, Reid J, Pocock M, Friedman CE, Mizikovsky D, Francois M, Palpant NJ, Needham EJ, Peralta M, Monte-Nieto GD, Jones LK, Smyth IM, Mehdiabadi NR, Bolk F, Janbandhu V, Yao E, Harvey RP, Chong JJH, Elliott DA, Stanley EG, Wiszniak S, Schwarz Q, James DE, Mills RJ, Porrello ER, Hudson JE. Vascular cells improve functionality of human cardiac organoids. Cell Rep. 2023 May 30;42(5):112322. doi: 10.1016/j.celrep.2023.112322. Epub 2023 Apr 26. PMID: 37105170.

21. Radisic M, Park H, Martens TP, Salazar-Lazaro JE, Geng W, Wang Y, Langer R, Freed LE, Vunjak-Novakovic G. Pre-treatment of synthetic elastomeric scaffolds by cardiac fibroblasts improves engineered heart tissue. J Biomed Mater Res A. 2008 Sep;86(3):713–24. doi: 10.1002/jbm.a.31578. PMID: 18041719; PMCID: PMC2775086.

22. Liau B, Christoforou N, Leong KW, Bursac N. Pluripotent stem cell-derived cardiac tissue patch with advanced structure and function. Biomaterials. 2011 Dec;32(35):9180–7. doi: 10.1016/j.biomaterials.2011.08.050. Epub 2011 Sep 8. PMID: 21906802; PMCID: PMC3190071.

23. Matsuura K, Masuda S, Haraguchi Y, Yasuda N, Shimizu T, Hagiwara N, Zandstra PW, Okano T. Creation of mouse embryonic stem cell-derived cardiac cell sheets. Biomaterials. 2011 Oct;32(30):7355–62. doi: 10.1016/j.biomaterials.2011.05.042. Epub 2011 Jul 31. PMID: 21807408.

24. Saini H, Navaei A, Van Putten A, Nikkhah M. 3D cardiac microtissues encapsulated with the co-culture of cardiomyocytes and cardiac fibroblasts. Adv Healthc Mater. 2015 Sep 16;4(13):1961–71. doi: 10.1002/adhm.201500331. Epub 2015 Jun 30. PMID: 26129820.

25. Iwamiya T, Matsuura K, Masuda S, Shimizu T, Okano T. Cardiac fibroblast-derived VCAM-1 enhances cardiomyocyte proliferation for fabrication of bioengineered cardiac tissue. Regen Ther. 2016 Jun 1;4:92–102. doi: 10.1016/j.reth.2016.01.005. PMID: 31245492; PMCID: PMC6581822.

26. Navaei A, Truong D, Heffernan J, Cutts J, Brafman D, Sirianni RW, Vernon B, Nikkhah M. PNIPAAm-based biohybrid injectable hydrogel for cardiac tissue engineering. Acta Biomater. 2016 Mar 1;32:10–23. doi: 10.1016/j.actbio.2015.12.019. Epub 2015 Dec 12. PMID: 26689467.

27. Ronaldson-Bouchard K, Ma SP, Yeager K, Chen T, Song L, Sirabella D, Morikawa K, Teles D, Yazawa M, Vunjak-Novakovic G. Advanced maturation of human cardiac tissue grown from pluripotent stem cells. Nature. 2018 Apr;556(7700):239-243. doi: 10.1038/s41586-018-0016-3. Epub 2018 Apr 4. Erratum in: Nature. 2019 Aug;572(7769):E16–E17. PMID: 29618819; PMCID: PMC5895513.

28. Beauchamp P, Jackson CB, Ozhathil LC, Agarkova I, Galindo CL, Sawyer DB, Suter TM, Zuppinger C. 3D Co-culture of hiPSC-Derived Cardiomyocytes With Cardiac Fibroblasts Improves Tissue-Like Features of Cardiac Spheroids. Front Mol Biosci. 2020 Feb 14;7:14. doi: 10.3389/fmolb.2020.00014. PMID: 32118040; PMCID: PMC7033479.

29. Fan D, Takawale A, Lee J, Kassiri Z. Cardiac fibroblasts, fibrosis and extracellular matrix remodeling in heart disease. Fibrogenesis Tissue Repair. 2012 Sep 3;5(1):15. doi: 10.1186/1755-1536-5-15. PMID: 22943504; PMCID: PMC3464725.

30. Hall C, Gehmlich K, Denning C, Pavlovic D. Complex Relationship Between Cardiac Fibroblasts and Cardiomyocytes in Health and Disease. J Am Heart Assoc. 2021 Feb;10(5):e019338. doi: 10.1161/JAHA.120.019338. Epub 2021 Feb 15. PMID: 33586463; PMCID: PMC8174279.

31. Bliley JM, Vermeer MCSC, Duffy RM, Batalov I, Kramer D, Tashman JW, Shiwarski DJ, Lee A, Teplenin AS, Volkers L, Coffin B, Hoes MF, Kalmykov A, Palchesko RN, Sun Y, Jongbloed JDH, Bomer N, de Boer RA, Suurmeijer AJH, Pijnappels DA, Bolling MC, van der Meer P, Feinberg AW. Dynamic loading of human engineered heart tissue enhances contractile function and drives a desmosome-linked disease phenotype. Sci Transl Med. 2021 Jul 21;13(603):eabd1817. doi: 10.1126/scitranslmed.abd1817. PMID: 34290054.

32. Zhu W, Zhao M, Mattapally S, Chen S, Zhang J. CCND2 Overexpression Enhances the Regenerative Potency of Human Induced Pluripotent Stem Cell-Derived Cardiomyocytes: Remuscularization of Injured Ventricle. Circ Res. 2018 Jan 5;122(1):88–96. doi: 10.1161/CIRCRESAHA.117.311504. Epub 2017 Oct 10. PMID: 29018036; PMCID: PMC5756126.

33. Zhao M, Nakada Y, Wei Y, Bian W, Chu Y, Borovjagin AV, Xie M, Zhu W, Nguyen T, Zhou Y, Serpooshan V, Walcott GP, Zhang J. Cyclin D2 Overexpression Enhances the Efficacy of Human Induced Pluripotent Stem Cell-Derived Cardiomyocytes for Myocardial Repair in a Swine Model of Myocardial Infarction. Circulation. 2021 Jul 20;144(3):210–228. doi: 10.1161/CIRCULATIONAHA.120.049497. Epub 2021 May 6. PMID: 33951921; PMCID: PMC8292228.

34 Kluesner MG, Nedveck DA, Lahr WS, Garbe JR, Abrahante JE, Webber BR, Moriarity BS. EditR: A Method to Quantify Base Editing from Sanger Sequencing. CRISPR J. 2018 Jun;1(3):239–250. doi: 10.1089/crispr.2018.0014. PMID: 31021262; PMCID: PMC6694769. Kluesner MG, Nedveck DA, Lahr WS, Garbe JR, Abrahante JE, Webber BR, Moriarity BS. EditR: A Method to Quantify Base Editing from Sanger Sequencing. CRISPR J. 2018 Jun;1(3):239-250. doi: 10.1089/crispr.2018.0014. PMID: 31021262; PMCID: PMC6694769.

35. Tyser RCV, Srinivas S. The First Heartbeat-Origin of Cardiac Contractile Activity. Cold Spring Harb Perspect Biol. 2020 Jul 1;12(7):a037135. doi: 10.1101/cshperspect.a037135. PMID: 31767652; PMCID: PMC7328461.

36. Männer J. When Does the Human Embryonic Heart Start Beating? A Review of Contemporary and Historical Sources of Knowledge about the Onset of Blood Circulation in Man. J Cardiovasc Dev Dis. 2022 Jun 9;9(6):187. doi: 10.3390/jcdd9060187. PMID: 35735816; PMCID: PMC9225347.

37. Spudich JA. Three perspectives on the molecular basis of hypercontractility caused by hypertrophic cardiomyopathy mutations. Pflugers Arch. 2019 May;471(5):701–717. doi: 10.1007/s00424-019-02259-2. Epub 2019 Feb 15. PMID: 30767072; PMCID: PMC6475635.

38. Ho CY, Carlsen C, Thune JJ, Havndrup O, Bundgaard H, Farrohi F, Rivero J, Cirino AL, Andersen PS, Christiansen M, Maron BJ, Orav EJ, Køber L. Echocardiographic strain imaging to assess early and late consequences of sarcomere mutations in hypertrophic cardiomyopathy. Circ Cardiovasc Genet. 2009 Aug;2(4):314–21. doi: 10.1161/CIRCGENETICS.109.862128. Epub 2009 Jun 19. PMID: 20031602; PMCID: PMC2773504.

39. Ho CY, López B, Coelho-Filho OR, Lakdawala NK, Cirino AL, Jarolim P, Kwong R, González A, Colan SD, Seidman JG, Díez J, Seidman CE. Myocardial fibrosis as an early manifestation of hypertrophic cardiomyopathy. N Engl J Med. 2010 Aug 5;363(6):552–63. doi: 10.1056/NEJMoa1002659. PMID: 20818890; PMCID: PMC3049917.

40. Ho CY, Sweitzer NK, McDonough B, Maron BJ, Casey SA, Seidman JG, Seidman CE, Solomon SD. Assessment of diastolic function with Doppler tissue imaging to predict genotype in preclinical hypertrophic cardiomyopathy. Circulation. 2002 Jun 25;105(25):2992–7. doi: 10.1161/01.cir.0000019070.70491.6d. PMID: 12081993.

41. Serraino GF, Jiritano F, Costa D, Ielapi N, Napolitano D, Mastroroberto P, Bracale UM, Andreucci M, Serra R. Metalloproteinases and Hypertrophic Cardiomyopathy: A Systematic Review. Biomolecules. 2023 Apr 11;13(4):665. doi: 10.3390/biom13040665. PMID: 37189412; PMCID: PMC10136246.

42. Luu RJ, Hoefler BC, Gard AL, Ritenour CR, Rogers MT, Kim ES, Coppeta JR, Cain BP, Isenberg BC, Azizgolshani H, Fajardo-Ramirez OR, García-Cardeña G, Lech MP, Tomlinson L, Charest JL, Williams C. Fibroblast activation in response to TGFβ1 is modulated by co-culture with endothelial cells in a vascular organ-on-chip platform. Front Mol Biosci. 2023 Jul 28;10:1160851. doi: 10.3389/fmolb.2023.1160851. PMID: 37577751; PMCID: PMC10421749.

43. Midgley AC, Rogers M, Hallett MB, Clayton A, Bowen T, Phillips AO, Steadman R. Transforming growth factor-β1 (TGF-β1)-stimulated fibroblast to myofibroblast differentiation is mediated by hyaluronan (HA)-facilitated epidermal growth factor receptor (EGFR) and CD44 co-localization in lipid rafts. J Biol Chem. 2013 May 24;288(21):14824–38. doi: 10.1074/jbc.M113.451336. Epub 2013 Apr 15. PMID: 23589287; PMCID: PMC3663506.

44. Aird EJ, Tompkins KJ, Ramirez MP, Gordon WR. Enhanced Molecular Tension Sensor Based on Bioluminescence Resonance Energy Transfer (BRET). ACS Sens. 2020 Jan 24;5(1):34–39. doi: 10.1021/acssensors.9b00796. Epub 2020 Jan 8. PMID: 31872754; PMCID: PMC7550199.

45. Ramirez MP, Anderson MJM, Kelly MD, Sundby LJ, Hagerty AR, Wenthe SJ, Odde DJ, Ervasti JM, Gordon WR. Dystrophin missense mutations alter focal adhesion tension and mechanotransduction. Proc Natl Acad Sci U S A. 2022 Jun 21;119(25):e2205536119. doi: 10.1073/pnas.2205536119. Epub 2022 Jun 14. PMID: 35700360; PMCID: PMC9231619.

46. Narikawa, M., Umemura, M., Tanaka, R. et al. Acute Hyperthermia Inhibits TGF-β1-induced Cardiac Fibroblast Activation via Suppression of Akt Signaling. Sci Rep 8, 6277 (2018). doi: 10.1038/s41598-018-24749-6.

47. Vallée A, Lecarpentier Y. TGF-β in fibrosis by acting as a conductor for contractile properties of myofibroblasts. Cell Biosci. 2019 Dec 9;9:98. doi: 10.1186/s13578-019-0362-3. PMID: 31827764; PMCID: PMC6902440.

48. Khalil H, Kanisicak O, Prasad V, Correll RN, Fu X, Schips T, Vagnozzi RJ, Liu R, Huynh T, Lee SJ, Karch J, Molkentin JD. Fibroblast-specific TGF-β-Smad2/3 signaling underlies cardiac fibrosis. J Clin Invest. 2017 Oct 2;127(10):3770–3783. doi: 10.1172/JCI94753. Epub 2017 Sep 11. PMID: 28891814; PMCID: PMC5617658.

49. Factor SM, Butany J, Sole MJ, Wigle ED, Williams WC, Rojkind M. Pathologic fibrosis and matrix connective tissue in the subaortic myocardium of patients with hypertrophic cardiomyopathy. J Am Coll Cardiol. 1991 May;17(6):1343–51. doi: 10.1016/s0735-1097(10)80145-7. PMID: 2016452.

50. Larson A, Codden CJ, Huggins GS, Rastegar H, Chen FY, Maron BJ, Rowin EJ, Maron MS, Chin MT. Altered intercellular communication and extracellular matrix signaling as a potential disease mechanism in human hypertrophic cardiomyopathy. Sci Rep. 2022 Mar 25;12(1):5211. doi: 10.1038/s41598-022-08561-x. PMID: 35338173; PMCID: PMC8956620.

51. Frantz S, Falcao-Pires I, Balligand JL, Bauersachs J, Brutsaert D, Ciccarelli M, Dawson D, de Windt LJ, Giacca M, Hamdani N, Hilfiker-Kleiner D, Hirsch E, Leite-Moreira A, Mayr M, Thum T, Tocchetti CG, van der Velden J, Varricchi G, Heymans S. The innate immune system in chronic cardiomyopathy: a European Society of Cardiology (ESC) scientific statement from the Working Group on Myocardial Function of the ESC. Eur J Heart Fail. 2018 Mar;20(3):445–459. doi: 10.1002/ejhf.1138. Epub 2018 Jan 15. PMID: 29333691; PMCID: PMC5993315.

52. Yu H, Gu L, Du L, Dong Z, Li Z, Yu M, Yin Y, Wang Y, Yu L, Ma H. Identification and analysis of key hypoxia- and immune-related genes in hypertrophic cardiomyopathy. Biol Res. 2023 Aug 9;56(1):45. doi: 10.1186/s40659-023-00451-4. PMID: 37559135; PMCID: PMC10410988.

53. Frieler RA, Mortensen RM. Immune cell and other noncardiomyocyte regulation of cardiac hypertrophy and remodeling. Circulation. 2015 Mar 17;131(11):1019–30. doi: 10.1161/CIRCULATIONAHA.114.008788. PMID: 25779542; PMCID: PMC4367123.

54. Sewanan LR, Schwan J, Kluger J, Park J, Jacoby DL, Qyang Y, Campbell SG. Extracellular Matrix From Hypertrophic Myocardium Provokes Impaired Twitch Dynamics in Healthy Cardiomyocytes. JACC Basic Transl Sci. 2019 Jul 24;4(4):495–505. doi: 10.1016/j.jacbts.2019.03.004. PMID: 31468004; PMCID: PMC6712054.

55. Wang X, Lu Y, Xie Y, Shen J, Xiang M. Emerging roles of proteoglycans in cardiac remodeling. Int J Cardiol. 2019 Mar 1;278:192–198. doi: 10.1016/j.ijcard.2018.11.125. Epub 2018 Nov 28. PMID: 30528626.

56. Alexanian M, Przytycki PF, Micheletti R, Padmanabhan A, Ye L, Travers JG, Gonzalez-Teran B, Silva AC, Duan Q, Ranade SS, Felix F, Linares-Saldana R, Li L, Lee CY, Sadagopan N, Pelonero A, Huang Y, Andreoletti G, Jain R, McKinsey TA, Rosenfeld MG, Gifford CA, Pollard KS, Haldar SM, Srivastava D. A transcriptional switch governs fibroblast activation in heart disease. Nature. 2021 Jul;595(7867):438–443. doi: 10.1038/s41586-021-03674-1. Epub 2021 Jun 23. PMID: 34163071; PMCID: PMC8341289.

57. Nagaraju CK, Dries E, Popovic N, Singh AA, Haemers P, Roderick HL, Claus P, Sipido KR, Driesen RB. Global fibroblast activation throughout the left ventricle but localized fibrosis after myocardial infarction. Sci Rep. 2017 Sep 7;7(1):10801. doi: 10.1038/s41598-017-09790-1. PMID: 28883544; PMCID: PMC5589875.

58. Wang L, Wang Y, Su Y, Wang J, Xiao M, Xi X-Y, Chen B-X, Zhang Y, Dong Z, Zhao S, Yang M-F. Activation of cardiac fibroblasts identifies high-risk hypertrophic cardiomyopathy. Journal of Nuclear Medicine Jun 2022, 63 (supplement 2) 2295.

59. Schlittler M, Pramstaller PP, Rossini A, De Bortoli M. Myocardial Fibrosis in Hypertrophic Cardiomyopathy: A Perspective from Fibroblasts. Int J Mol Sci. 2023 Oct 2;24(19):14845. doi: 10.3390/ijms241914845. PMID: 37834293; PMCID: PMC10573356.

60. Kim JB, Porreca GJ, Song L, Greenway SC, Gorham JM, Church GM, Seidman CE, Seidman JG. Polony multiplex analysis of gene expression (PMAGE) in mouse hypertrophic cardiomyopathy. Science. 2007 Jun 8;316(5830):1481-4. doi: 10.1126/science.1137325. PMID: 17556586.

61. Frangogiannis N. Transforming growth factor-β in tissue fibrosis. J Exp Med. 2020 Feb 13;217(3):e20190103. doi: 10.1084/jem.20190103. PMID: 32997468; PMCID: PMC7062524.

62. Dobaczewski M, Chen W, Frangogiannis NG. Transforming growth factor (TGF)-β signaling in cardiac remodeling. J Mol Cell Cardiol. 2011 Oct;51(4):600–6. doi: 10.1016/j.yjmcc.2010.10.033. Epub 2010 Nov 6. PMID: 21059352; PMCID: PMC3072437.

63. Frangogiannis NG. Cardiac fibrosis. Cardiovasc Res. 2021 May 25;117(6):1450–1488. doi: 10.1093/cvr/cvaa324. PMID: 33135058; PMCID: PMC8152700.

64. Teekakirikul P, Eminaga S, Toka O, Alcalai R, Wang L, Wakimoto H, Nayor M, Konno T, Gorham JM, Wolf CM, Kim JB, Schmitt JP, Molkentin JD, Norris RA, Tager AM, Hoffman SR, Markwald RR, Seidman CE, Seidman JG. Cardiac fibrosis in mice with hypertrophic cardiomyopathy is mediated by non-myocyte proliferation and requires Tgf-β. J Clin Invest. 2010 Oct;120(10):3520–9. doi: 10.1172/JCI42028. Epub 2010 Sep 1. PMID: 20811150; PMCID: PMC2947222.

65. Tyska MJ, Hayes E, Giewat M, Seidman CE, Seidman JG, Warshaw DM. Single-molecule mechanics of R403Q cardiac myosin isolated from the mouse model of familial hypertrophic cardiomyopathy. Circ Res. 2000 Apr 14;86(7):737–44. doi: 10.1161/01.res.86.7.737. PMID: 10764406.

66. Debold EP, Schmitt JP, Patlak JB, Beck SE, Moore JR, Seidman JG, Seidman C, Warshaw DM. Hypertrophic and dilated cardiomyopathy mutations differentially affect the molecular force generation of mouse alpha-cardiac myosin in the laser trap assay. Am J Physiol Heart Circ Physiol. 2007 Jul;293(1):H284–91. doi: 10.1152/ajpheart.00128.2007. Epub 2007 Mar 9. PMID: 17351073.

67. Koitabashi N, Danner T, Zaiman AL, Pinto YM, Rowell J, Mankowski J, Zhang D, Nakamura T, Takimoto E, Kass DA. Pivotal role of cardiomyocyte TGF-β signaling in the murine pathological response to sustained pressure overload. J Clin Invest. 2011 Jun;121(6):2301–12. doi: 10.1172/JCI44824. Epub 2011 May 2. PMID: 21537080; PMCID: PMC3104748.

68. Alamo L, Ware JS, Pinto A, Gillilan RE, Seidman JG, Seidman CE, Padrón R. Effects of myosin variants on interacting-heads motif explain distinct hypertrophic and dilated cardiomyopathy phenotypes. Elife. 2017 Jun 13;6:e24634. doi: 10.7554/eLife.24634. PMID: 28606303; PMCID: PMC5469618.

69. Verma BK, Chatterjee A, Kondaiah P, Gundiah N. Substrate Stiffness Modulates TGF-β Activation and ECM-Associated Gene Expression in Fibroblasts. Bioengineering. 2023; 10(9):998. doi: 10.3390/bioengineering10090998.

70. Felisbino MB, Rubino M, Travers JG, Schuetze KB, Lemieux ME, Anseth KS, Aguado BA, McKinsey TA. Substrate stiffness modulates cardiac fibroblast activation, senescence, and proinflammatory secretory phenotype. Am J Physiol Heart Circ Physiol. 2024 Jan 1;326(1):H61–H73. doi: 10.1152/ajpheart.00483.2023. Epub 2023 Oct 27. PMID: 37889253.

71. Achterberg VF, Buscemi L, Diekmann H, Smith-Clerc J, Schwengler H, Meister JJ, Wenck H, Gallinat S, Hinz B. The nano-scale mechanical properties of the extracellular matrix regulate dermal fibroblast function. J Invest Dermatol. 2014 Jul;134(7):1862–1872. doi: 10.1038/jid.2014.90. Epub 2014 Feb 13. PMID: 24670384.

72. Wang H, Haeger SM, Kloxin AM, Leinwand LA, Anseth KS. Redirecting valvular myofibroblasts into dormant fibroblasts through light-mediated reduction in substrate modulus. PLoS One. 2012;7(7):e39969. doi: 10.1371/journal.pone.0039969. Epub 2012 Jul 13. PMID: 22808079; PMCID: PMC3396623.

73. Zampieri M, Berteotti M, Ferrantini C, Tassetti L, Gabriele M, Tomberli B, Castelli G, Cappelli F, Stefàno P, Marchionni N, Coppini R, Olivotto I. Pathophysiology and Treatment of Hypertrophic Cardiomyopathy: New Perspectives. Curr Heart Fail Rep. 2021 Aug;18(4):169–179. doi: 10.1007/s11897-021-00523-0. Epub 2021 Jun 20. PMID: 34148184.

74. Gersh BJ, Maron BJ, Bonow RO, Dearani JA, Fifer MA, Link MS, Naidu SS, Nishimura RA, Ommen SR, Rakowski H, Seidman CE, Towbin JA, Udelson JE, Yancy CW; American College of Cardiology Foundation/American Heart Association Task Force on Practice Guidelines; American Association for Thoracic Surgery; American Society of Echocardiography; American Society of Nuclear Cardiology; Heart Failure Society of America; Heart Rhythm Society; Society for Cardiovascular Angiography and Interventions; Society of Thoracic Surgeons. 2011 ACCF/AHA guideline for the diagnosis and treatment of hypertrophic cardiomyopathy: executive summary: a report of the American College of Cardiology Foundation/American Heart Association Task Force on Practice Guidelines. Circulation. 2011 Dec 13;124(24):2761–96. doi: 10.1161/CIR.0b013e318223e230. Epub 2011 Nov 8. PMID: 22068435.

75. Kim SH, Kim SO, Han S, Hwang KW, Lee CW, Nam GB, Choi KJ, Kim DH, Song JM, Kang DH, Song JK, Kim CH, Kim YH. Long-term comparison of apical versus asymmetric hypertrophic cardiomyopathy. Int Heart J. 2013;54(4):207–11. doi: 10.1536/ihj.54.207. PMID: 23924932.

76. Sherrid MV, Pearle G, Gunsburg DZ. Mechanism of benefit of negative inotropes in obstructive hypertrophic cardiomyopathy. Circulation. 1998 Jan 6-13;97(1):41–7. doi: 10.1161/01.cir.97.1.41. Erratum in: Circulation 1998 Mar 17;97(10):1026. PMID: 9443430.

77. Verlinden NJ, Coons JC. Disopyramide for Hypertrophic Cardiomyopathy: A Pragmatic Reappraisal of an Old Drug. Pharmacotherapy. 2015 Dec;35(12):1164–72. doi: 10.1002/phar.1664. PMID: 26684556.

78. Palandri C, Santini L, Argirò A, Margara F, Doste R, Bueno-Orovio A, Olivotto I, Coppini R. Pharmacological Management of Hypertrophic Cardiomyopathy: From Bench to Bedside. Drugs. 2022 Jun;82(8):889–912. doi: 10.1007/s40265-022-01728-w. Epub 2022 Jun 13. PMID: 35696053; PMCID: PMC9209358.

79. Stewart S, Mason DT, Braunwald E. Impaired rate of left ventricular filling in idiopathic hypertrophic subaortic stenosis and valvular aortic stenosis. Circulation. 1968 Jan;37(1):8–14. doi: 10.1161/01.cir.37.1.8. PMID: 5688694.

80. Klein MD, Lane FJ, Gorlin R. Effect of left ventricular size and shape upon the hemodynamics of subaortic stenosis. Am J Cardiol. 1965;15:773–781. doi: 10.1016/0002-9149(65)90379-6. PMID: 14299372.

81. Wilson WS, Criley JM, Ross RS. Dynamics of left ventricular emptying in hypertrophic subaortic stenosis. A cineangiographic and hemodynamic study. Am Heart J. 1967 Jan;73(1):4–16. doi: 10.1016/0002-8703(67)90303-1. PMID: 6066683.

82. Spudich JA. Hypertrophic and dilated cardiomyopathy: four decades of basic research on muscle lead to potential therapeutic approaches to these devastating genetic diseases. Biophys J. 2014 Mar 18;106(6):1236–49. doi: 10.1016/j.bpj.2014.02.011. PMID: 24655499; PMCID: PMC3985504.

83. Ashrafian H, Redwood C, Blair E, Watkins H. Hypertrophic cardiomyopathy: a paradigm for myocardial energy depletion. Trends Genet. 2003 May;19(5):263–8. doi: 10.1016/S0168-9525(03)00081-7. PMID: 12711218.

84. Ommen SR, Ho CY, Asif IM, Balaji S, Burke MA, Day SM, Dearani JA, Epps KC, Evanovich L, Ferrari VA, Joglar JA, Khan SS, Kim JJ, Kittleson MM, Krittanawong C, Martinez MW, Mital S, Naidu SS, Saberi S, Semsarian C, Times S, Waldman CB. 2024 AHA/ACC/AMSSM/HRS/PACES/SCMR Guideline for the Management of Hypertrophic Cardiomyopathy: A Report of the American Heart Association/American College of Cardiology Joint Committee on Clinical Practice Guidelines. Circulation. 2024 Jun 4;149(23):e1239–e1311. doi: 10.1161/CIR.0000000000001250. Epub 2024 May 8. PMID: 38718139.

85. Monda E, Bakalakos A, Rubino M, Verrillo F, Diana G, De Michele G, Altobelli I, Lioncino M, Perna A, Falco L, Palmiero G, Elliott PM, Limongelli G. Targeted Therapies in Pediatric and Adult Patients With Hypertrophic Heart Disease: From Molecular Pathophysiology to Personalized Medicine. Circ Heart Fail. 2023 Aug;16(8):e010687. doi: 10.1161/CIRCHEARTFAILURE.123.010687. Epub 2023 Jul 21. PMID: 37477018.

86. Ho CY, Day SM, Axelsson A, Russell MW, Zahka K, Lever HM, Pereira AC, Colan SD, Margossian R, Murphy AM, Canter C, Bach RG, Wheeler MT, Rossano JW, Owens AT, Bundgaard H, Benson L, Mestroni L, Taylor MRG, Patel AR, Wilmot I, Thrush P, Vargas JD, Soslow JH, Becker JR, Seidman CE, Lakdawala NK, Cirino AL; VANISH Investigators; Burns KM, McMurray JJV, MacRae CA, Solomon SD, Orav EJ, Braunwald E. Valsartan in early-stage hypertrophic cardiomyopathy: a randomized phase 2 trial. Nat Med. 2021 Oct;27(10):1818-1824. doi: 10.1038/s41591-021-01505-4. Epub 2021 Sep 23. PMID: 34556856; PMCID: PMC8666141.

87. Argirò A, Zampieri M, Marchi A, Cappelli F, Del Franco A, Mazzoni C, Cecchi F, Olivotto I. Stage-specific therapy for hypertrophic cardiomyopathy. Eur Heart J Suppl. 2023 Apr 26;25(Suppl C):C155–C161. doi: 10.1093/eurheartjsupp/suad042. PMID: 37125313; PMCID: PMC10132571.

88. Zhang J, Fan G, Zhao H, Wang Z, Li F, Zhang P, Zhang J, Wang X, Wang W. Targeted inhibition of Focal Adhesion Kinase Attenuates Cardiac Fibrosis and Preserves Heart Function in Adverse Cardiac Remodeling. Sci Rep. 2017 Feb 22;7:43146. doi: 10.1038/srep43146. PMID: 28225063; PMCID: PMC5320468.

89. Gilbert CJ, Longenecker JZ, Accornero F. ERK1/2: An Integrator of Signals That Alters Cardiac Homeostasis and Growth. Biology (Basel). 2021 Apr 20;10(4):346. doi: 10.3390/biology10040346. PMID: 33923899; PMCID: PMC8072600.

90. Wu X, Simpson J, Hong JH, Kim KH, Thavarajah NK, Backx PH, Neel BG, Araki T. MEK-ERK pathway modulation ameliorates disease phenotypes in a mouse model of Noonan syndrome associated with the Raf1(L613V) mutation. J Clin Invest. 2011 Mar;121(3):1009–25. doi: 10.1172/JCI44929. Epub 2011 Feb 21. PMID: 21339642; PMCID: PMC3049402.

