## Supplementary Figure 1 for "HCM-associated mutations in MYH6/7 drive pathologic expression of TGF-β1 in cardiomyocytes within weeks of developmental specification"

Supplementary Figure 1. EHT compaction over time.

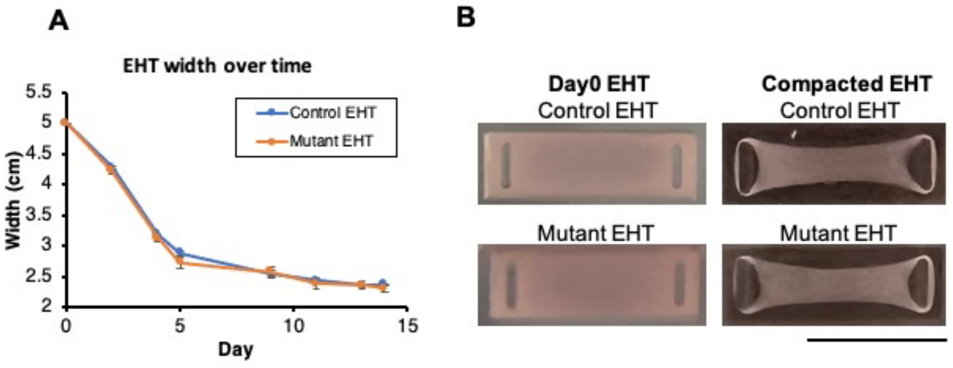


(A) EHT width change after fabrication.

(A) Representative images for control and mutant EHTs at Day 0 and Day 12 after fabrication. (Scale bar = 1 cm) (n = 6 Bright-field images for each condition)
