## Supplementary Figure 2 for "HCM-associated mutations in MYH6/7 drive pathologic expression of TGF-β1 in cardiomyocytes within weeks of developmental specification"

Supplementary Figure 2. Alignment and distribution of hiPSC-CMs of EHTs

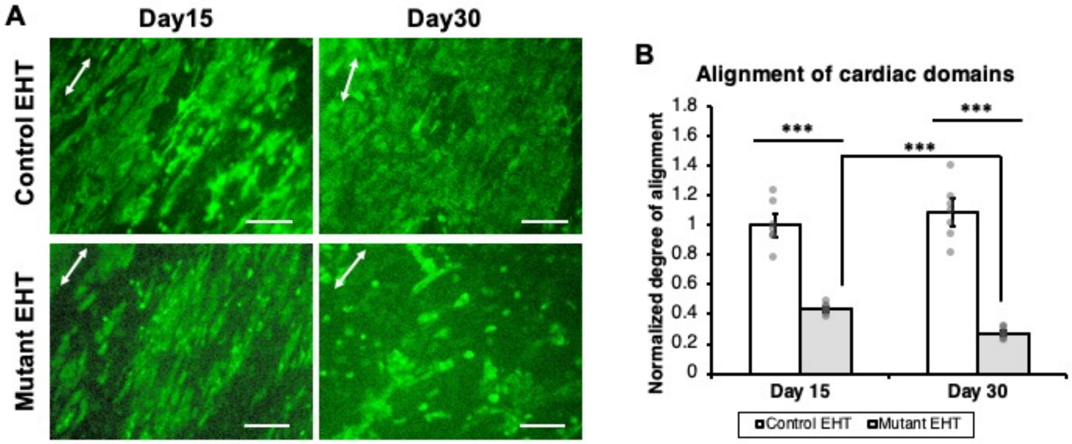


(A) Green contrast corresponds to calcium-sensitive dye, Fluo-4 AM. Images show the maximal fluorescent signal during calcium cycling in cardiomyocytes.

(B) Quantitative analysis of hiPSC-CM alignment based on contrast corresponding to Fluo-4 AM fluorescence emission. (*** $p < 0.005$; Student $t$ test; n = 6 immunostained images for each condition at each time point)
