## Supplementary Figure 3 for "HCM-associated mutations in MYH6/7 drive pathologic expression of TGF-β1 in cardiomyocytes within weeks of developmental specification"

Supplementary Figure 3. Force of contraction of wild type and mutant EHTs over time and with electrical pacing.

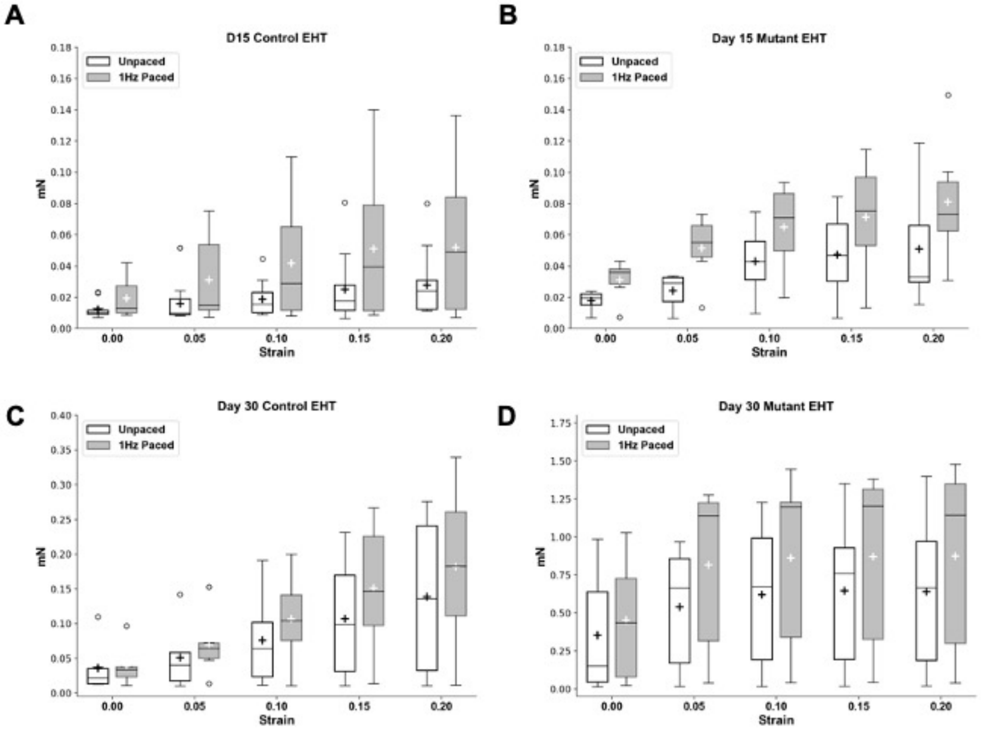


(A) control EHT at Day 15 after fabrication. (Three EHTs for each condition and two experimental replicates, n = 6 for both mutant and control groups)

(B) mutant EHT at Day 15 after fabrication. (Three EHTs for each condition and two experimental replicates, n = 6 for both mutant and control groups)

(C) control EHT at Day 30 after fabrication. (Three EHTs for each condition and two experimental replicates, n = 6 for both mutant and control groups)

(D) mutant EHT at Day 30 after fabrication. (Three EHTs for each condition and two experimental replicates, n = 6 for both mutant and control groups)

During the force generation measurement, each EHT was gradually stretched to 120% of the original length, contraction forces under 1Hz field stimulated or unpaced conditions were assessed.
