## Supplementary Figure 4 for "HCM-associated mutations in MYH6/7 drive pathologic expression of TGF-β1 in cardiomyocytes within weeks of developmental specification"

Supplementary Figure 4. Western blot of pFAK and FAK for control and mutant EHTs at Day 15 and Day 30.

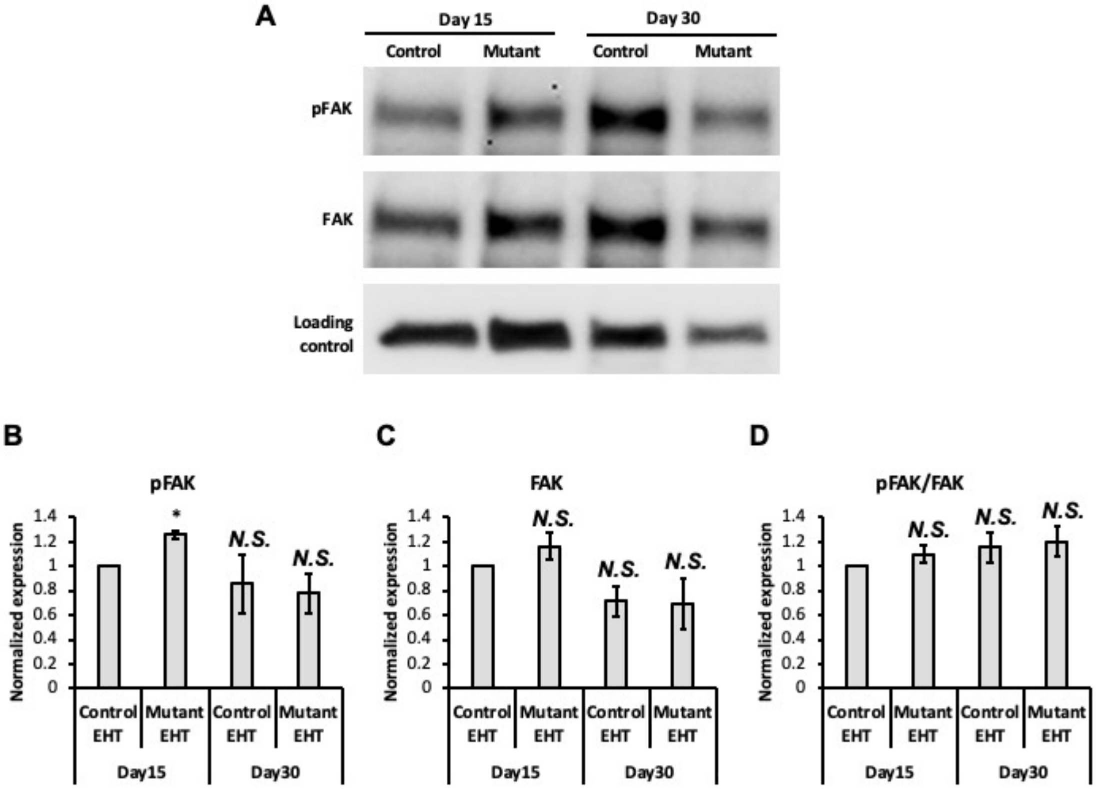


(A) Image of Western blot for pFAK and FAK with GAPDH as the loading control for Day 15 samples and $\alpha$-tubulin for Day 30 samples.

(B) Quantitative result of Western blot for pFAK expression (normalized to Day 15 Control EHT). (*N.S.*: not significant; *$p < 0.05$; Student $t$ test; one EHT for each condition and two experimental replicates, n = 2 for both control and mutant groups)

(C) Quantitative result of Western blot for FAK expression (normalized to Day 15 Control EHT). (*N.S.*: not significant; Student $t$ test; one EHT for each condition and two experimental replicates, n = 2 for both control and mutant groups)

(D) Quantitative result of Western blot for pFAK/FAK (normalized to Day 15 Control EHT). (*N.S.*: not significant; Student $t$ test; one EHT for each condition and two experimental replicates, n = 2 for both control and mutant groups)

Note that the results of the Western blot for pFAK and FAK in both control and mutant EHTs showed a decrease when comparing Day 30 to Day 15. This could be due to the increased expression of overall proteins at Day 30, resulting in a reduced proportion of pFAK and FAK. Therefore, while the same amount of total protein was loaded for Western blot, the levels of pFAK and FAK at Day 30 appeared lower than those at Day 15.
