## Supplementary Figure 5 for "HCM-associated mutations in MYH6/7 drive pathologic expression of TGF-β1 in cardiomyocytes within weeks of developmental specification"

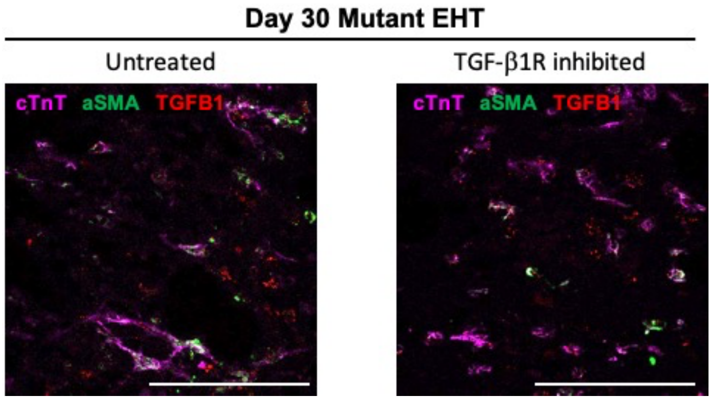


Representative images of cTnT, αSMA, and TGF-β1-stained sections for Day 30 mutant EHT with or without TGF-β1R inhibition. (Scale bar = 100 μm)
