## Supplementary Figure 6 for "HCM-associated mutations in MYH6/7 drive pathologic expression of TGF-β1 in cardiomyocytes within weeks of developmental specification"

Supplementary Figure 6. TGF-β1 mRNA expression in fibroblast and activated fibroblast.

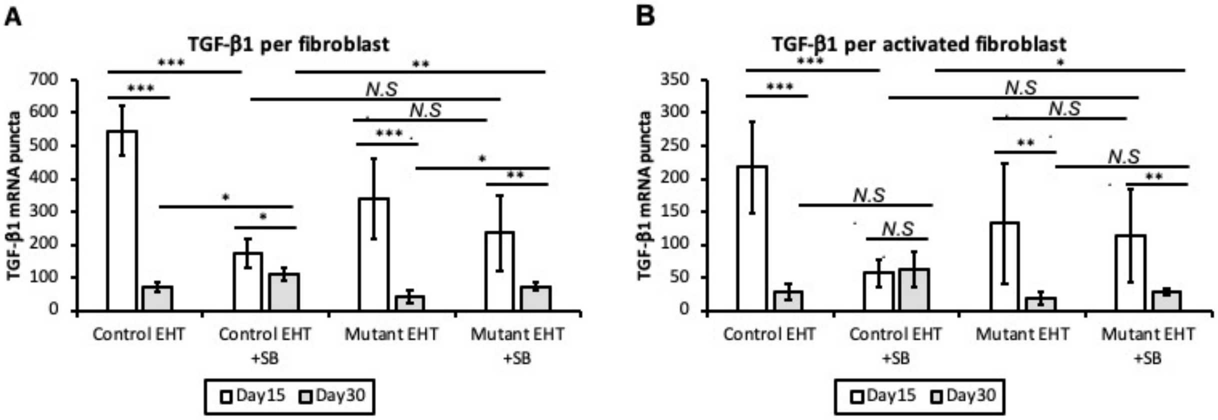


(A) Quantitative analysis of TGF-β1 mRNA expression per fibroblast in both control and mutant EHTs at Day 15 and 30 with or without inhibition. (***$p < 0.005$; **$p < 0.01$; *$p < 0.05$; *N.S.*: not significant; Student *t* test; n = 6 FISH-IHC stained images for each condition)

(B) Quantitative analysis of TGF-β1 mRNA expression per activated fibroblast in both control and mutant EHTs at Day 15 and 30 with or without inhibition. (***$p < 0.005$; **$p < 0.01$; *$p < 0.05$; *N.S.*: not significant; Student *t* test; n = 6 FISH-IHC stained images for each condition)
