## Supplementary Table 1 for "HCM-associated mutations in MYH6/7 drive pathologic expression of TGF-β1 in cardiomyocytes within weeks of developmental specification"

Supplementary Table 1. List of antibodies used in this study

| # | Antibody | Manufacturer, Catalog # | Host, Isotype | Classification | Dilution |
|---|---|---|---|---|---|
| 1 | Cardiac Troponin T (13-11) | Epredia, MS-295-P1 | Mouse IgG | Monoclonal | 1:200 (IHC) |
| 2 | $\alpha$-Smooth Muscle Actin | NOVUSBIO, NBP1-30894 | Rabbit IgG | Polyclonal | 1:200 (IHC) |
| 3 | Fibroblasts Antibody (TE-7) | NOVUSBIO, NBP2-50082 | Mouse IgG1 | Monoclonal | 1:200 (IHC) |
| 4 | Phospho-FAK (Tyr397) | Invitrogen, 700255 | Rabbit IgG | Recombinant monoclonal | 1:200 (IHC) 1:500 (WB) |
| 5 | FAK (34Q36) | Invitrogen, AHO1272 | Mouse IgG2b, $\kappa$ | Monoclonal | 1:200 (IHC) 1:500 (WB) |
| 6 | Collagen I | Invitrogen, PA5-95137 | Rabbit IgG | Polyclonal | 1:200 (IHC) |
| 7 | Fibronectin | Invitrogen, PA5-29578 | Rabbit IgG | Polyclonal | 1:200 (IHC) |
| 8 | Alexa Fluor 488 goat anti-mouse IgG | Invitrogen, A11001 | Goat IgG | Polyclonal | 1:500 (IHC) |
| 9 | Alexa Fluor 647 goat anti-mouse IgG | Invitrogen, A12136 | Goat IgG | Polyclonal | 1:500 (IHC) |
| 10 | Alexa Fluor 488 goat anti-rabbit IgG | Invitrogen, A11008 | Goat IgG | Polyclonal | 1:500 (IHC) |
| 11 | Alexa Fluor 647 goat anti-rabbit IgG | Invitrogen, A21244 | Goat IgG | Polyclonal | 1:500 (IHC) |
| 12 | GAPDH (D16H11) | Cell Signaling Technology, #5174 | Rabbit IgG | Recombinant monoclonal | 1:1000 (WB) |
| 13 | $\alpha$-Tubulin | Cell Signaling Technology, #2144 | Rabbit IgG | Polyclonal | 1:1000 (WB) |
| 14 | Anti-rabbit IgG, HRP-linked antibody | Cell Signaling Technology, #7074 | Goat | Polyclonal | 1:3000 (WB) |
| 15 | Anti-mouse IgG, HRP-linked antibody | Cell Signaling Technology, #7076 | Horse | Polyclonal | 1:3000 (WB) |
